# Host Gut Motility and Bacterial Competition Drive Instability in a Model Intestinal Microbiota

**DOI:** 10.1101/052985

**Authors:** Travis J. Wiles, Matthew L. Jemielita, Ryan P. Baker, Brandon H. Schlomann, Savannah L. Logan, Julia Ganz, Ellie Melancon, Judith S. Eisen, Karen Guillemin, Raghuveer Parthasarathy

## Abstract

The gut microbiota is a complex consortium of microorganisms with the ability to influence important aspects of host health and development. Harnessing this ‘microbial organ’ for biomedical applications requires clarifying the degree to which host and bacterial factors act alone or in combination to govern the stability of specific lineages. To address this we combined bacteriological manipulation and light sheet fluorescence microscopy to monitor the dynamics of a defined two-species microbiota within the vertebrate gut. We observed that the interplay between each population and the gut environment produced distinct spatiotemporal patterns. Consequently, one species dominates while the other experiences dramatic collapses that are well fit by a stochastic mathematical model. Modeling revealed that bacterial competition could only partially explain the observed phenomena, suggesting that a host factor is also important in shaping the community. We hypothesized the host determinant to be gut motility, and tested this mechanism by measuring colonization in hosts with enteric nervous system dysfunction due to mutation in the Hirschsprung disease locus *ret*. In mutant hosts we found reduced gut motility and, confirming our hypothesis, robust coexistence of both bacterial species. This study provides evidence that host-mediated spatial structuring and stochastic perturbation of communities along with bacterial competition drives population dynamics within the gut. In addition, this work highlights the capacity of the enteric nervous system to affect stability of gut microbiota constituents, demonstrating that the ‘gut-brain axis’ is bidirectional. Ultimately, these findings will help inform disease mitigation strategies focused on engineering the intestinal ecosystem.

## INTRODUCTION

Trillions of microbial cells representing hundreds of species make up the human intestinal microbiota. This multispecies symbiont supports activities as diverse as host development, nutrient acquisition, immune system education, neural function, and defense against pathogens [1-5]. Changes in microbiota diversity and functional composition have been linked with a variety of human disorders, including obesity, colon cancer, opportunistic infection, and inflammatory bowel disease [6, 7]. A major goal of host-microbe systems biology is to clarify the ecological factors that determine microbiota integrity by meshing experimental techniques and quantitative modeling. Insights derived from such efforts will inspire the design of novel therapeutic strategies for microbiota-associated diseases.

An unresolved question is whether the host and microbiota function independently or together to govern the dynamics and stability of individual bacterial lineages. Addressing this requires identifying the interactions that arise within the spatially complex and heterogeneous environment of the vertebrate gut. However, progress toward this goal has been hindered due to the technical limitations associated with directly observing intestinal communities. Typical interrogation of vertebrate intestinal microbiota involves phylogenetic profiling of fecal material using high-throughput sequencing of 16S ribosomal RNA (rRNA) genes. This technique is blind to the spatial structure of microbial communities, which is known in general to strongly influence interactions [8-10] and has recently been predicted to be important for microbiota stability [11]. Sequencing-based studies also have low sensitivity to temporal structure owing to both experimental and analytic constraints. Experimentally, metagenomic time-series data remain rare and cannot reach the sampling frequencies necessary to capture interactions occurring at the timescales of microbial division or intestinal flux. Analytically, sequencing data yield only relative, rather than absolute, taxonomic abundances, which severely confounds the inference of interaction networks [12, 13]. Furthermore, such methods to date have employed deterministic Lotka-Volterra models [12, 14, 15] that, even with noise terms representing measurement error, neglect the possibility of fundamentally stochastic or discontinuous interactions among constituents.

Because our knowledge of the factors that shape interactions within host-associated ecosystems is incomplete, contemporary theoretical models have been forced to rely on assumptions that may not realistically mirror microbe-microbe and host-microbe relationships. For example, biochemical and physical inputs from the animal host that likely act on microbial constituents are often ignored [16]. It is important to unravel the extent to which microbiota integrity is simply an intrinsic property of the microbes, which could be recapitulated *in vitro* with co-culture experiments or *in silico* with bacterial metabolic network models, or an emergent property of the host-microbe system. Developing accurate accounts of the ecological interactions that manifest within the gut will require model systems that enable manipulation of microbial colonization and measurement methods that can characterize microbial populations *in vivo* with high spatial and temporal resolution [17].

Toward this end, we employ here larval zebrafish as a model vertebrate host coupled with light sheet fluorescence microscopy (LSFM) [18] as a minimally invasive interrogation method to examine population dynamics within a defined gut microbiota. Zebrafish larvae are highly amenable to gnotobiotic techniques and can be reared germ-free (GF) in large numbers [19]. At four days post-fertilization (dpf) larvae possess an open and functional digestive tract that is permissive to microbial colonization, the timing of which is controlled by adding bacteria to the water column. Importantly, larval zebrafish share many physiological traits with humans, including aspects of innate immunity, neurological development, and intestinal function [20]. Therefore, interactions between zebrafish and their microbial symbionts are expected to reflect analogous interactions that occur in other vertebrates. LSFM, combined with the optical transparency of larval zebrafish, enables three-dimensional visualization of the entire intestine with single-bacterium resolution, rapid image acquisition to avoid blurring due to intestinal motility, and extended live imaging with low phototoxicity [21, 22]. This experimental setup provides an unprecedented opportunity to investigate ecological interactions within the vertebrate intestine at a range of spatial and temporal scales.

With this model system we found that an apparent competitive interaction between two species native to the zebrafish gut, *Aeromonas veronii* and *Vibrio cholerae*, is characterized by sudden and catastrophic collapses of the *Aeromonas* population, which appear to be driven by mechanical forces related to host intestinal motility. The differential behavior of these two species can be explained by their distinct biogeography and community architecture within the intestine. Further mapping of *Aeromonas*-*Vibrio* dynamics motivated a quantitative stochastic model with parameters that could be independently derived from traditional abundance measurements and image-based time-series analysis. Ultimately, this model allowed us to predict the consequences of altering the host environment through genetic disruption of the enteric nervous system (ENS) via a mutation in *ret*, a gene locus associated with human Hirschsprung disease (OMIM 164761), which stabilized the *Aeromonas* population and neutralized competition with *Vibrio*. This work reveals a synergy between bacterial competition and host-mediated spatial structuring of microbiota in determining population dynamics and stability—a feature that is likely mirrored in more complex host-microbe systems such as the human gut.

## RESULTS

### *Aeromonas* and *Vibrio* exhibit an apparent competitive interaction in the zebrafish gut

The intestinal microbiota of larval zebrafish is dominated by bacterial lineages belonging to the Gammaproteobacteria [23, 24]. In a prior investigation we found that two representative isolates native to the zebrafish intestinal tract, *Aeromonas veronii* strain ZOR0001, hereafter referred to as *Aeromonas*, and a *Vibrio* strain, ZWU0020, similar in 16S rRNA gene sequence to *Vibrio cholerae* and hereafter referred to as *Vibrio*, exhibit an apparent negative interaction in GF larval zebrafish, with populations of *Aeromonas* that were several orders of magnitude lower in di-associations with *Vibrio* than in mono-associations [25]. Interestingly, only a modest suppression was observed in *in vitro* competition experiments [25]. To begin to untangle the importance of host and bacterial factors in facilitating the *in vivo* interaction between *Aeromonas* and *Vibrio*, we used a succession assay in which GF larval zebrafish were first colonized by *Aeromonas* to high abundance, and then challenged by invading populations of *Vibrio* (Fig 1A).

**Fig 1.**
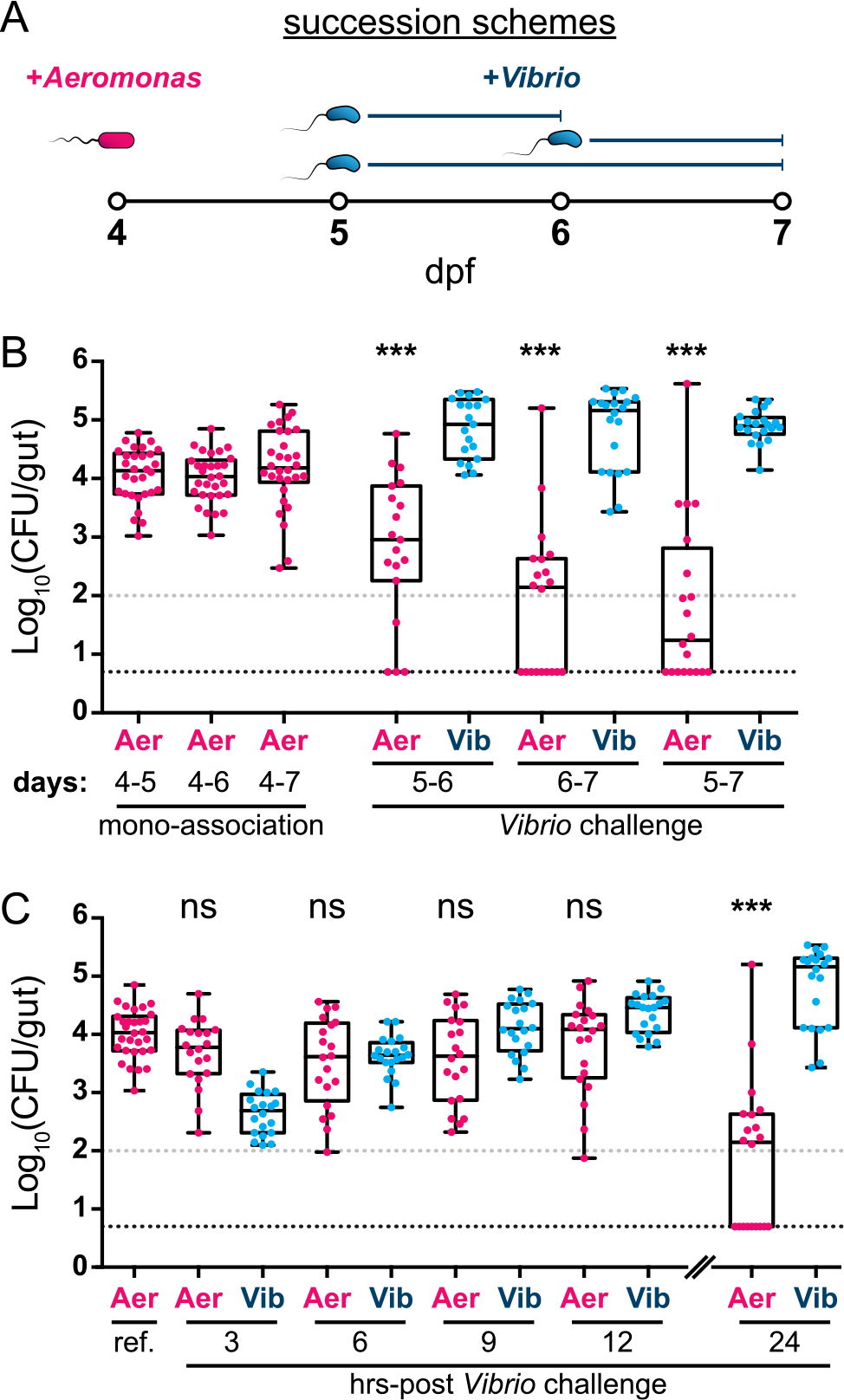
***Aeromonas* and *Vibrio* exhibit an apparent competitive interaction within the larval zebrafish intestine**. (A) Graphical overview of succession schemes used to characterize *Aeromonas*-*Vibrio* interactions. *Aeromonas* is allowed to colonize GF larvae at 4 dpf followed by addition of *Vibrio* to the water column at 5 or 6 dpf for 24 or 48 hours prior to enumeration of abundances by dissection and serial plating techniques. (B, left) *Aeromonas* abundances after different mono-association durations and (B, right) *Aeromonas* and *Vibrio* abundances after different *Vibrio* challenge periods. Statistical significance of *Aeromonas* abundances after *Vibrio* challenge compared to respective mono-association reference populations (i.e. ‘5-6’ vs. ‘4-6’; ‘6-7’ vs. ‘4-7’; ‘5-7’ vs. ‘4-7’) was determined by an unpaired t-test. (C) Time course analysis of *Aeromonas* and *Vibrio* abundances determined by dissection and plating at three-hour intervals over a 12-hour period starting at 6 dpf. Additionally plotted are an *Aeromonas* mono-association reference population and 24 hour *Aeromonas* and *Vibrio* populations previously plotted in 1B (’4-6’ and ‘6-7’, respectively). Statistical significance of *Aeromonas* abundances to the mono-association reference population (ref.) was determined by an unpaired t-test. CFU=colony-forming units; ***=p<0.0001; ns=not significant; N>19/condition. Gray and black dashed lines in panels B and C denote limits of quantification and detection, respectively.

We first enumerated total bacterial abundance by gut dissection and standard plating techniques. In mono-associations beginning at 4 dpf and extending 24, 48, or 72 hours *Aeromonas* populations consistently reach 10^4^ colony-forming units (CFU) per host (Fig 1B). In contrast, challenge of established *Aeromonas* populations with *Vibrio* over various 24 or 48 hour temporal windows leads to dramatically lower *Aeromonas* abundance as well as increased host-to-host variation and frequent extinction events (Fig 1B). Under these conditions *Vibrio* exhibits only modest deviations in abundance compared to mono-association (S1A and S1B Fig). Of note, *Aeromonas* populations are not destabilized upon self-challenge by newly introduced *Aeromonas*, and *Vibrio* does not induce collapses in established *Vibrio* populations (S1C and S1D Fig). These results indicate that subsequent waves of colonizing bacteria alone do not account for the observed competitive interaction and that it is inter-specific in nature. We also verified that competition between *Aeromonas* and *Vibrio* is dependent on the host environment, as there was not an appreciable difference between their abundances during *in vitro* mono- and co-culture experiments (S1E Fig). To determine if the abundance of *Vibrio* correlates with a reduction of *Aeromonas* populations *in vivo*, we performed a time course experiment in which zebrafish were sacrificed and assayed every 3 hours for 12 hours after inoculation with *Vibrio*. We found that *Vibrio* rapidly infiltrates *Aeromonas*-colonized intestines and steadily increases in number over the 12-hour assay period (Fig 1C). Surprisingly, we did not detect a concomitant decline in *Aeromonas*, implying that *Aeromonas* populations do not merely respond proportionally to the abundance of *Vibrio* (Fig 1C).

### *Aeromonas* population dynamics are altered during Vibrio challenge

To further explore the interactions driving *Aeromonas-Vibrio* competition, we turned to LSFM. Imaging fluorescently marked variants of each species during mono-association revealed that they have noticeably different intestinal biogeographies and behavior (Fig 2). Populations of *Vibrio* largely comprise planktonic, highly motile cells that appear capable of sampling all available regions within the intestine (S1, S2, and S3 Movie). Quantifying the bacterial density along the anterior-posterior axis (Methods), we find that *Vibrio* is most abundant in the anterior bulb (Fig 2B and 2C), with an overabundance within several micrometers of the epithelial wall that may be the result of hydrodynamic interactions (S2 Fig) [26]. In contrast, *Aeromonas* is most abundant in the midgut and largely takes the form of dense, non-motile clusters with a smaller subpopulation of motile individuals (Fig 2D and 2E; S4 and S5 Movie).

**Fig 2.**
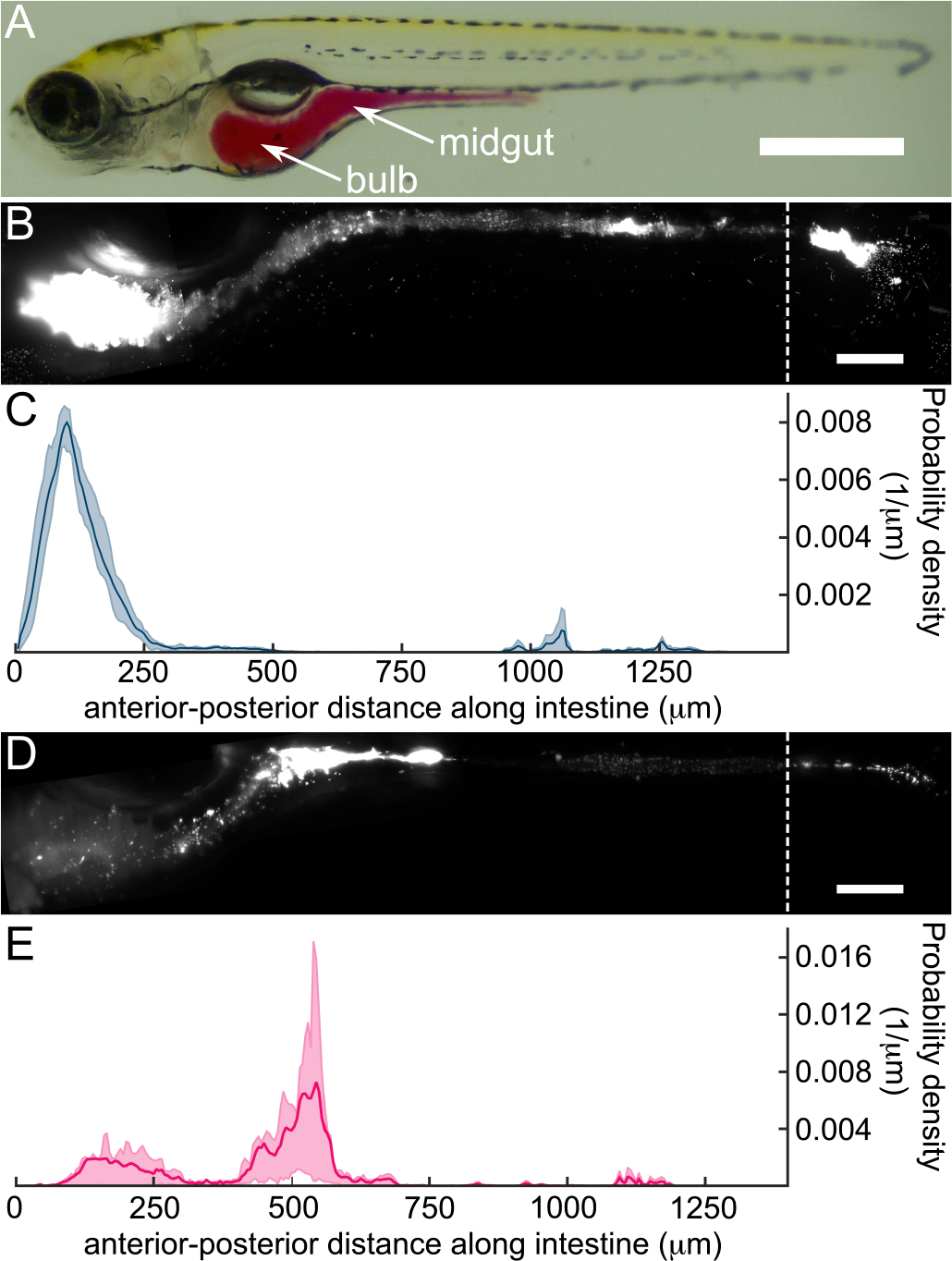
***Vibrio* and *Aeromonas* have distinct community architectures and biogeographies within the larval zebrafish intestine**. (A) A larval zebrafish at 5 dpf; the intestine is highlighted by phenol red dye via microgavage [61]. Scale bar: 500 µm. (B) A maximum intensity projection (MIP) of *Vibrio* in the larval intestine. Scale bar: 100 µm. (C) The probability density of *Vibrio* along the intestinal axis. From (B) and (C), we see that *Vibrio* is predominantly localized in the anterior bulb. (D) MIP of *Aeromonas* in the larval intestine. Scale bar: 100 µm. (E) The probability density of *Aeromonas* along the intestinal axis. (D) and (E) show that *Aeromonas* is predominantly localized in the midgut, with a smaller population in the anterior bulb.

To identify the temporal dynamics of the two-member community, we performed *in vivo* live imaging experiments using LSFM starting approximately 2 hours following the challenge of established *Aeromonas* populations with *Vibrio*. Three-dimensional images spanning the intestine were obtained for each species for durations of roughly 12-15 hours at 20-minute intervals, which is shorter than each species’ approximate one-hour doubling time (Methods). Fig 3A shows maximum intensity projections of *Aeromonas* and *Vibrio* in a representative larval zebrafish intestine at three time points spanning a four-hour interval (S6 Movie). The abundance of each species over several hours is plotted in Fig 3B and 3C for two fish, representative of twelve fish that we examined. Similar population abundance plots for all zebrafish are provided in S3 Fig. We found that *Vibrio* populations smoothly expand and exhibit a growth rate of 0.8 ± 0.3 hr.^-1^ (mean ± std. dev.), similar to that derived from plating data (0.60 ± 0.22 hr.^-1^, Fig 1C). Strikingly, growth of *Aeromonas* populations is sporadically interrupted by dramatic collapse events, dropping in abundance by multiple orders of magnitude within an hour (Fig 3A, 3B, and 3C).

**Fig 3.**
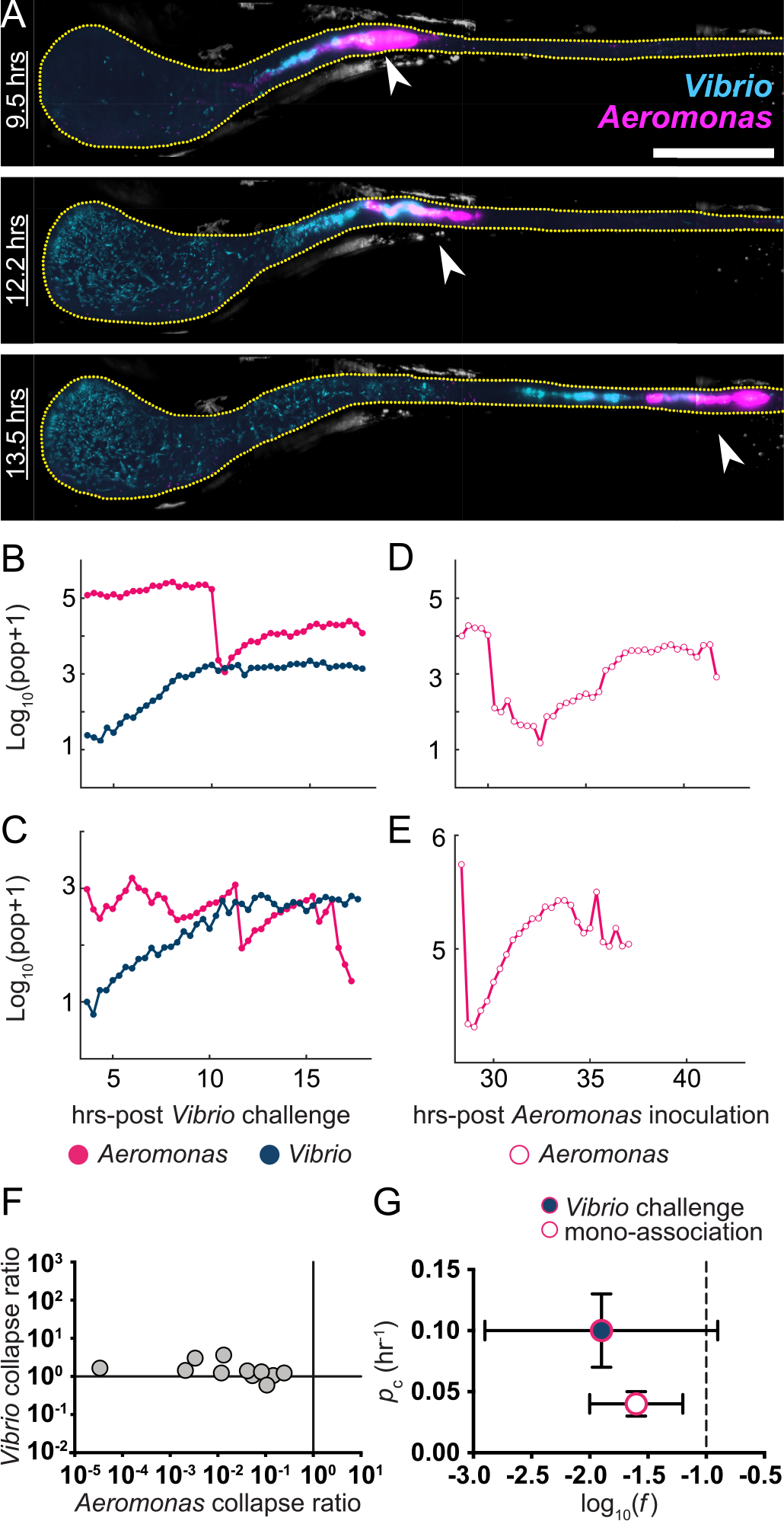
***Aeromonas* populations experience collapse events during *Vibrio* challenge and mono-association**. (A) MIPs of *Aeromonas* (magenta) and *Vibrio* (cyan) in a larval zebrafish intestine. Scale bar: 200 µm. The fish was initially colonized at 4 dpf with *Aeromonas*, challenged 24 hours later by inoculation with *Vibrio*, and then imaged every 20 minutes for 14 hours. The times indicated denote hours post-challenge. In all images, the region shown spans about 80% of the intestine, with the anterior on the left. Image contrast in both color channels is enhanced for clarity. Yellow dotted line roughly indicates the lumenal boundary. As time progresses, the anterior growth of *Vibrio* as well as abrupt changes in the *Aeromonas* distribution (arrows) are evident. (B,C) Total bacterial abundance, derived from image data, for *Aeromonas* and *Vibrio* in two representative fish inoculated and challenged as in panel A, as a function of time following the *Vibrio* inoculation. Sharp drops of over an order of magnitude in the *Aeromonas* population, but not the *Vibrio* population, are evident. (D,E) Total abundance for *Aeromonas* in mono-associations as a function of time post-inoculation, in two representative fish. Collapses are also observed, though in general the populations recover to approximately pre-collapse levels. (F) The ratio, *f*, of the post-collapse to the pre-collapse population for *Aeromonas* challenged by *Vibrio*, which span many orders of magnitude (horizontal axis). At the same time points, the *Vibrio* populations are essentially unchanged, with ratios of post- to pre-collapse populations close to one (vertical axis). (G) Characteristics of *Aeromonas* population collapses. Circles and bars indicate the mean and standard deviation, respectively, of *f* and *p*_c_, the rate of collapse occurrence, for both mono-associations and *Aeromonas* challenged by *Vibrio*. The dashed line at *f =* 0.1 indicates the threshold for identification of collapses.

To determine if *Aeromonas* collapses occur in the absence of *Vibrio*, we examined live imaging data of *Aeromonas* mono-associations over a similar time frame. We detected clear instances of *Aeromonas* population collapses under these conditions (Fig 3D and 3E). However, in contrast to *Vibrio*-associated collapses, *Aeromonas* was found to consistently recover from collapses during mono-association. Additionally, *Aeromonas* collapse events associated with mono-association were smaller and more uniform. Defining collapses as events in which the population decreases by at least a factor of ten within one hour, and assigning their magnitude *f* as the ratio of the population after collapse to that before, we find that for *Aeromonas* challenged by *Vibrio*, log_10_(*f*) = -1.9 ± 1.0 (mean ± std. dev.) (Fig 3F and 3G). The ratio of the *Vibrio* population before and after *Aeromonas* collapse events within the same fish is approximately 1 (Fig 3F), corroborating observations from imaging and plating data that *Vibrio* is resistant to the perturbations that affect *Aeromonas*. We found that during *Aeromonas* mono-associations the magnitude of collapse events was about half of that observed in the presence of *Vibrio*, log_10_(*f*) = -1.6 ± 0.4 (Fig 3G). We also detected a greater rate of *Aeromonas* collapses during *Vibrio* challenge. Estimating the collapse probability per unit time, *p*_c_, as the total number of collapses in all fish divided by the total observation time, we find *p*_c_ = 0.10 ± 0.03 hr.^-1^ during *Vibrio* challenge and *p*_c_ = 0.04 ± 0.01 hr.^-1^ during *Aeromonas* mono-associations (Fig 3G), where the uncertainties are estimated by assuming an underlying Poisson process.

### *Aeromonas* and *Vibrio* are differentially resistant to intestinal motility

We next inspected the spatial structure and dynamics of each species to uncover clues regarding possible factors driving *Aeromonas* collapses. A conspicuous feature of the larval zebrafish intestine, like most animal intestines, is that it undergoes periodic contractions that propagate along its length. We found that *Vibrio* populations, being made up of motile, planktonic individuals, are almost completely unaffected by the mechanics of intestinal motility (Fig 4A). Like a liquid filling its container, populations of *Vibrio* quickly adapt to the contracting and expanding space with surprisingly little change in their distribution (Fig 4B; S3 Movie). In contrast, the rigid and largely non-motile clusters of *Aeromonas*, localized to the narrow midgut, are strongly affected by intestinal contractions (Fig 4C and 4D; S7 Movie). These observations support the hypothesis that forces exerted on this two-member community by intestinal motility give rise to rare and stochastic expulsion of *Aeromonas* while leaving *Vibrio* unperturbed. Corroborating this notion, we indeed observe posterior transport of *Aeromonas* in collapse events during live imaging experiments (Fig 3A; S6 Movie).

**Fig 4.**
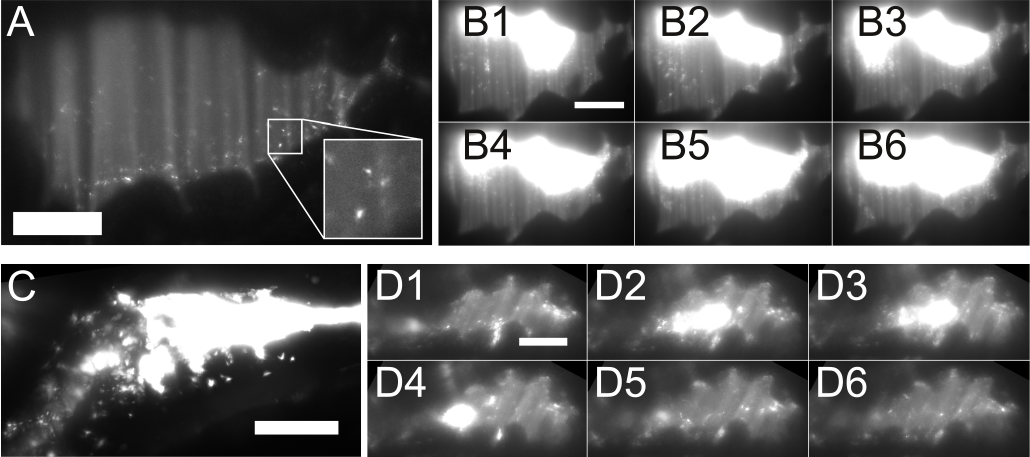
***Aeromonas* and *Vibrio* exhibit different dynamics within the zebrafish intestine**. (A) An optical section of *Vibrio* mono-associated with a larval zebrafish, showing the anterior bulb region. The population consists of discrete, highly motile individuals. (B) A montage of images from a time-series of *Vibrio* in the bulb, during which the overall distribution of the population remains stable. Time between frames: 1 second. (C) An optical section of the midgut of *Aeromonas* mono-associated with a larval zebrafish. Dense clusters are evident. (D) A montage of images from a time-series of *Aeromonas* in the midgut, during which the overall distribution of the population changes considerably. Time between frames: 1 second. Scale bars: 50 µm.

*Aeromonas* collapses occur with or without the presence of *Vibrio*, but these collapses have different significance in the two cases for the overall *Aeromonas* abundance. One can imagine several possible explanations for this. We first asked whether the growth rate of *Aeromonas* post-collapse is lower if *Vibrio* is present. The data show that this is not the case. Mono-associated *Aeromonas* have a post-collapse growth rate of 0.74 ± 0.1 hr.^-1^ (mean ± std. dev., *N*=5 collapses), whereas *Vibrio*-challenged *Aeromonas* have 0.64 ± 0.2 hr.^-1^ (*N*=4 collapses). We next asked whether the presence of *Vibrio* leads to changes in the mechanics of intestinal motility. To test this, we imaged intestinal motility in larval zebrafish using differential interference contrast microscopy (DIC) [27] and calculated the dominant period and amplitude of intestinal contractions. Comparing GF fish with *Vibrio* or *Aeromonas* mono-associated fish, or fish in which *Aeromonas* is challenged after 24 hours by *Vibrio*, there is no notable difference in period or amplitude (S4 Fig). The consequences of intestinal motility on *Aeromonas* collapse properties are clearly different in the mono-association and challenge cases, however, as indicated by changes in collapse magnitudes and rates (*f* and *p*_c_). We also note that during challenge experiments, the gross spatial distribution *of Vibrio* is similar to its distribution during mono-association, while there is considerable broadening in the spatial distribution of *Aeromonas* when challenged (S5 Fig). Finally, a conceptually minimal model of interaction is that with *Vibrio* present, the resources available to *Aeromonas* post-collapse are less than with *Vibrio* absent, thereby placing a limit on its potential for recovery. We assess this possibility quantitatively below by estimating carrying capacities, and we also examine the synergistic consequences of changes to the carrying capacity and collapse properties.

### Synergy between stochastic collapse events and competition with *Vibrio* underlies *Aeromonas* population dynamics

Thus far our data suggest that *Aeromonas* is susceptible to stochastic disturbances mediated by host intestinal motility, and that its recovery from these disturbances is altered in the presence of *Vibrio*. If this is the case, we should be able to build a quantitative model that reflects these data, explains the high variance observed in plating assays (Fig 1), and offers insights into the differential outcomes between mono-association and challenge experiments. The model we construct is illustrated schematically in Fig 5. Consider a bacterial species exhibiting logistic growth, with growth rate *r* and carrying capacity *K* (Fig 5A and 5B); in other words, the population *N* grows with time *t* according to:

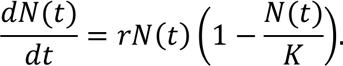

Superimposed on this are rare collapses, during which the population drops to *f* times its pre-collapse value, where *f* is between 0 and 1, and after which it resumes logistic growth (Fig 5C). The collapses are stochastic and modeled as Poisson processes; i.e. they occur at random with some probability per unit time *p*_c_ (Fig 5D). This model arises in many ecological contexts, and some of its mathematical properties have been explored in various studies [28]. Of course, this model incorporates stochastic population collapses by construction, and so does not predict them from first principles. However, the parameter values that emerge from fitting such a model to the data can, as shown below, yield quantitative insights into the mechanisms underlying the observed dynamics that are not evident from mere visual inspection of the raw data.

**Fig 5.**
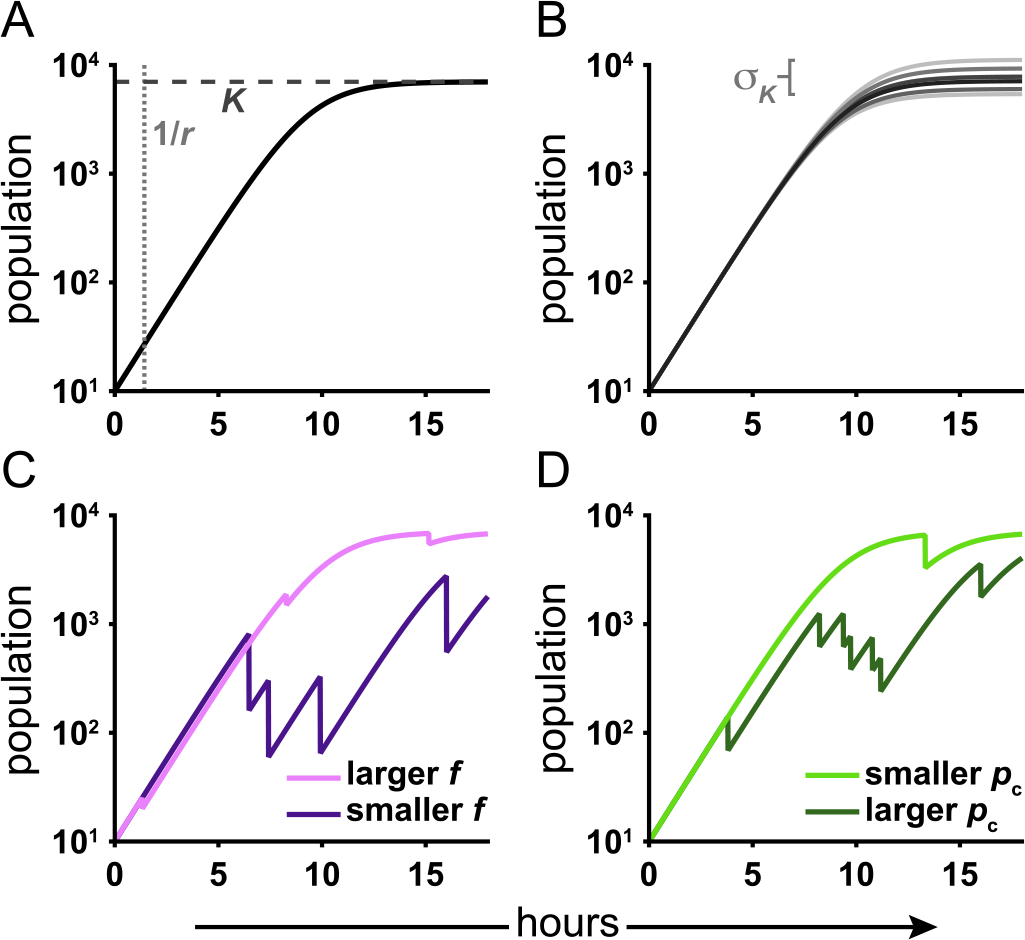
**Schematic of a model of growth punctuated by collapses**. (A) The model is based on simple logistic growth, which is characterized by two parameters, the growth rate, *r*, and carrying capacity, *K*. (B) We also include a parameter characterizing variability in the carrying capacity. Stochastic collapses are governed by two parameters: (C) the fraction of the population remaining after a collapse, *f*, and (D) the probability per unit time of a collapse, *p*_c_.

Simulating ensembles of populations that exhibit the above dynamics, we examine the mean and, importantly, the standard deviation of the population at discrete terminal time points, as these are statistics that allow direct comparison to results from plating assays. As shown in detail in the Supporting Text (S1 Text), the apparent dependence of the model on the parameters, *r, K, p*_c_, and *f*, collapses to two effective parameters. The growth rate, *r*, is both independently known and irrelevant for the conditions considered, and the dynamics depend on the combination *z =* -*p*_c_ log_10_(*f*) rather than on *p*_c_ and *f* independently. Values of the two remaining relevant parameters, *K* and *z*, which characterize the carrying capacity and the collapse dynamics, respectively, determine the model predictions for the mean and variance of populations. A grid search through the (*K, z*) space for the values that minimize the distance between the predicted and observed *Aeromonas* population statistics gives the best-fit model parameters. Additional details and discussion are provided in the Supporting Text (S1 Text). It is important to note that because our imaging data revealed that *Aeromonas* is often in a state of experiencing or recovering from collapse events, the observed population is likely never close to *K*, and thus we cannot simply use the mean of the bacterial abundance to estimate *K*. Rather, we must use a model to infer the carrying capacity that would yield the observed populations.

Using *Aeromonas* abundance data obtained by gut dissection and plating 24 hours post *Vibrio* challenge (Fig 1B, ‘6-7’), we find best-fit parameters log_10_(*K*) = 3.2 ± 0.5 and *z =* 0.13 ± 0.05 hr.^-1^, the latter providing a constraint on *p*_c_ and *f* together. We can *independently* estimate *p*_c_ and *f* from imaging-derived data (Fig 3). As noted previously, for *Aeromonas* challenged by *Vibrio*, we find *p*_c_ = 0.10 ± 0.03 hr.^-1^ and log_10_(*f*) = -1.9 ± 0.3 (mean ± std. error), yielding *z =* 0.19 ± 0.06 hr.^-1^, which is consistent with the plating-derived value. The agreement between the separately determined measures of *z* is remarkable, as it indicates that the statistical properties inferred from an ensemble of populations at a discrete time point are consistent with the properties inferred from the temporal dynamics within individual hosts. As expected, log_10_(*K*) is greater than the observed mean *Aeromonas* abundance at 24 hours, since the model-derived *K* represents an upper bound for the population in the absence of any stochastic collapses or *Vibrio* competition. As another test of consistency, we note that simulating our stochastic model for 48 hours post-challenge using the best-fit parameters determined from plating experiments 24 hours post-challenge predicts a mean and standard deviation of log_10_(population+1) of 1.3 ± 0.3 and 1.5 ± 0.2, respectively, in agreement with the observed plating values of 1.7 ± 0.3 and 1.6 ± 0.3 (Fig 1B). All of these assessments support the conclusion that the observed population dynamics are governed by a mechanism of stochastic collapse.

We can also apply this model to *Aeromonas* mono-association data. Here, the variance of the plating-derived populations is small (Fig 1B), likely due to comparatively rare and/or weak collapses as discussed earlier. For reasons described in detail in the Supporting Text, this hinders robust determination of *z*, though *K* remains well fit. We find that *z =* 0.01 ± 0.01 hr.^-1^ and log_10_(*K*) = 4.2 ± 0.1. From live imaging data, *p*_c_ = 0.04 ± 0.01 hr.^-1^ and log_10_(*f*) = -1.6 ± 0.2, from which *z =* 0.06 ± 0.02 hr.^-1^ (S1 Text). Our identification of thresholds is, by construction, only sensitive to collapses of a factor of 10 or more in magnitude (i.e. log_10_(*f*) ≤ -1), so our estimate of *f*, and therefore *z*, is biased toward larger values.

The above analysis yields insights into the nature of the competition between *Aeromonas* and *Vibrio* that are not obvious from simple visual inspection of the data. The carrying capacity (*K*) experienced by *Aeromonas*, as estimated by our model, is only one order of magnitude lower in the presence of *Vibrio* (log_10_(*K*) = 3.2 ± 0.5) than when *Vibrio* is absent (log_10_(*K*) = 4.2 ± 0.1). However, the observed abundance of *Aeromonas* is suppressed by more than two orders of magnitude: mean(log_10_(population+1)) = 1.7 ± 0.3 and 4.1 ± 0.1 when challenged by *Vibrio* and mono-associated, respectively (Fig 1B). These results suggest that the combined effect of stochastic collapses, which are likely driven by the host environment, and a reduced carrying capacity, as a result of competition with *Vibrio*, has a far greater influence on population dynamics than either mechanism would provide alone.

### Mutant hosts lacking enteric nervous system function stabilize *Aeromonas* in the face of *Vibrio* challenge

Together, our experimental data and quantitative predictions indicate that a synergy between competition with *Vibrio* and host-mediated stochastic disturbances underlies the destabilization of *Aeromonas* populations within the larval zebrafish intestine. Our model predicts that if the host factor intestinal motility were reduced, *Aeromonas* populations would be more stable despite the presence of *Vibrio*. To test this hypothesis, we carried out succession assays in mutant zebrafish hosts essentially lacking a functional enteric nervous system (ENS) because of disruption of the gene encoding the *Ret* tyrosine kinase, which is critical for ENS development [29]. Using DIC microscopy to assess intestinal motility, we found that *ret* mutant larvae (*ret*^-/-^) still exhibit rhythmic contractions, but with different characteristics than wild-type (*ret*^+/+^) and heterozygous siblings (*ret*^+/-^) (S8 and S9 Movie). Because we observed that *ret*^+/+^ and *ret*^+/-^ animals are phenotypically similarity with regard to gut motility and that the *ret1^hu2846^* mutant allele is recessive we further designate *ret*^+/+^ and *ret*^+/-^ as ‘wild type’. Computational analysis of time-series DIC images allows quantification of the displacement of intestinal tissue during contractile waves (Methods). The average peak amplitude of longitudinal contractions is greater in wild-type than in *ret* mutant larvae, and in both genotypes declines with age (Fig 6A). At 6 dpf a considerable fraction of *ret* mutant larvae show low amplitudes, similar to the quiescent state observed in both genotypes at 7 dpf (Fig 6A). Though the amplitude of intestinal contractions might not be directly related to the magnitude or rate of *Aeromonas* collapse events, it is reasonable to expect some monotonic relationship between the two, as they both reflect intestinal activity. Therefore, we would expect to observe stabilization of *Vibrio*-challenged *Aeromonas* populations in *ret* mutant hosts only during challenge periods starting at 6 dpf when the difference in intestinal motility between the genotypes is greatest. Indeed, *Vibrio* challenge of established *Aeromonas* populations between 5 and 6 dpf yielded the same decrease in *Aeromonas* abundance in both *ret* mutant hosts and wild types (Fig 6B). In contrast, *Aeromonas* populations were significantly stabilized during *Vibrio* challenge from 6 to 7 dpf in *ret* mutant hosts and in fact were statistically indistinguishable from a reference *Aeromonas* mono-association (Fig 6B). These results provide strong evidence that ENS-driven intestinal motility contributes to the shaping of this model two-member community by facilitating their apparent competitive interaction.

**Fig 6.**
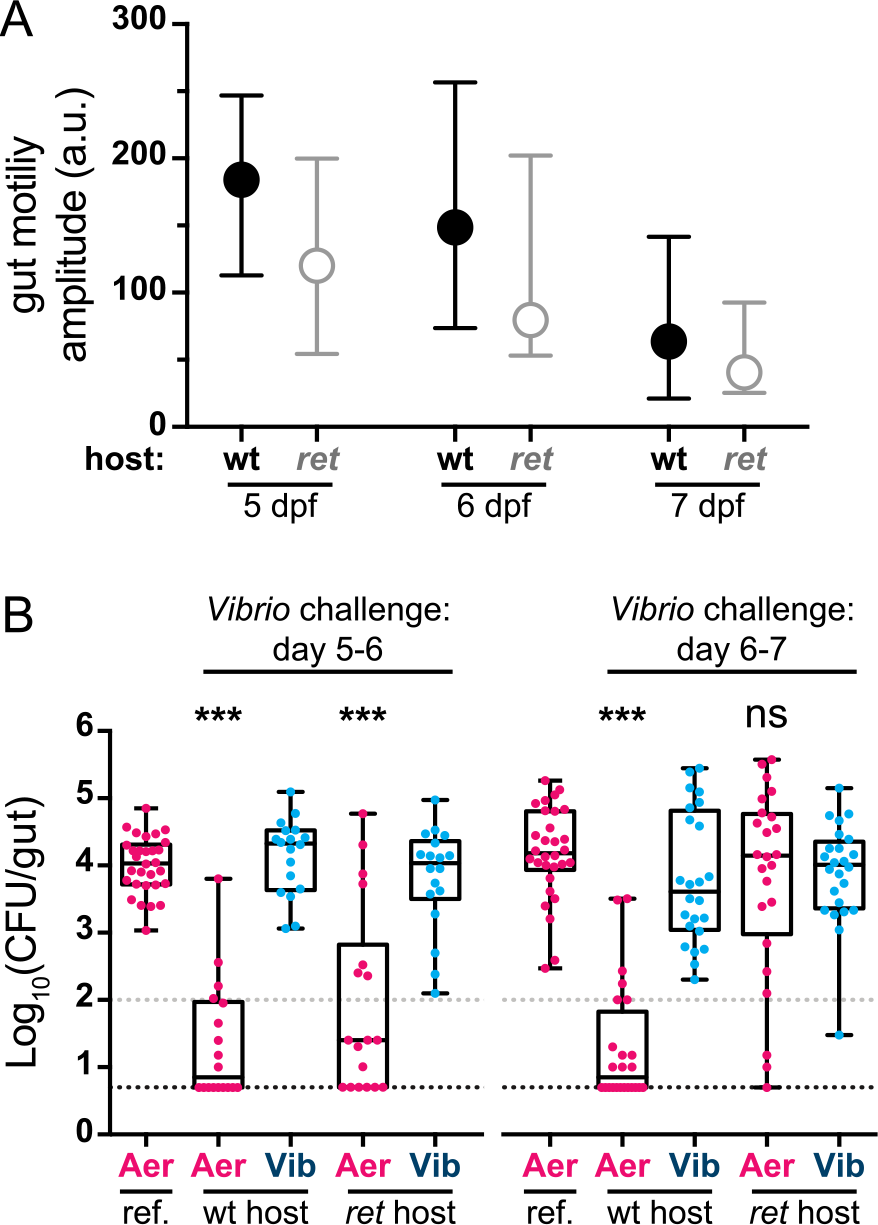
**Intestinal motility and bacterial competition are altered in *ret* mutant zebrafish hosts**. (A) Amplitudes of periodic contraction along the intestine for wild-type and *ret* mutant zebrafish at various ages. Circles indicate medians and bars indicate first and third quartiles. (B) GF wild-type and *ret* heterozygous hosts (wt) were raised together with *ret* homozygous mutant hosts (*ret*) and colonized at 4 dpf with *Aeromonas*. At 5 (left) or 6 (right) dpf *Vibrio* was added to the water column for 24 hours prior to whole gut dissection and serial plating to enumerate bacterial abundances. Additionally plotted are respective *Aeromonas* mono-association reference (ref.) populations from Fig 1B (left, ‘4-6’; right, ‘4-7’). The difference between *Aeromonas* abundance during challenge and mono-association was determined by an unpaired t-test. CFU=colony-forming units; ***=p<0.0001; ns=not significant; N>18/condition. Gray and black dashed lines denote limits of quantification and detection, respectively.

## DISCUSSION

A better understanding of the factors that influence the dynamics and stability of host-associated microbial communities would allow insights into their assembly [30-32], fluctuations during periods of normal health [12, 14, 15, 33, 34], and responses to perturbation [35, 36], as well as aid the development of diagnostic and treatment strategies for human diseases [17]. Building a working knowledge of these processes has been impeded by the technical difficulties associated with examining bacterial populations within their native host environments. In humans, the approach generally taken has been to infer inter-species interactions from coarsely sampled sequencing-based metagenomic time-series experiments performed on fecal samples [12-14, 34, 37]. However, such procedures largely disregard spatial information and generally assume particular functional forms for interactions (e.g. deterministic Lotka-Volterra dynamics [12, 14, 15]). Moreover, measurement noise and missing information about absolute abundances in metagenomic data place severe limits on the quantitative determination of interaction strengths, even if the models are accurate descriptors of the microbial systems [12]. Therefore, basic questions regarding inter-species competition in the intestine, particularly the extent to which it is determined by the microbes themselves, properties of the host environment, or a combination of the two, remain largely unanswered.

For these reasons we set out to investigate bacterial population dynamics within the vertebrate intestine using a combination of absolute abundance measurements, time-series imaging, and quantitative modeling. Though our system is minimal, consisting of two bacterial species and a larval zebrafish host, it has revealed factors we expect to be of broad relevance to other animal-associated microbiota. Most notably, in this model system the emergence of the apparent competition between *Aeromonas* and *Vibrio* is driven in large part by the physical activity of the host, namely the motility of the intestine. In mutant zebrafish hosts that have reduced intestinal motility due to mutation of the gene *ret*, which impairs ENS development and function, competition between these bacterial species is offset (Fig 6). Motility and mass transport are, of course, key attributes of animal intestinal tracts. The finding that the mechanical nature of the host environment has a major role in shaping bacterial communities suggests that models of microbiota based on *in vitro* competition assays or modeling of metabolic networks [16, 38] will, by themselves, be insufficient for predicting and accurately describing community structure and dynamics. This is in line with the recent observation that dietary alteration of intestinal transit in a murine model can lead to compositional shifts in the gut microbiota [39]. Moreover, it provides a mechanism by which host genotype can influence community composition. Corroborating this notion, human patients with Hirschsprung disease, which is a gastrointestinal motility disorder commonly associated with mutation of *ret*, have been found to harbor dysbiotic microbial communities [40, 41].

The differential susceptibility of our two model bacterial species to intestinal motility can be explained by their distinct community architectures. Highly motile *Vibrio* are relatively unaffected by intestinal contractions, which is in contrast to the large, non-motile aggregates of *Aeromonas* (Fig 4). Earlier observations of a related *A. veronii* strain showed higher growth rates for aggregated bacteria compared to planktonic [21], suggesting a tradeoff between enhanced growth and resistance to population level perturbations. In general, we suspect that the spatial structure of microbial communities within the intestine will be an important determinant of their dynamics and a key consideration for the generation of successful predictive models.

We are able to construct a quantitative model of the observed *Aeromonas* dynamics that consists of growth punctuated by stochastic collapses. Data derived from gut dissection, in which many fish are sampled at a single time point, can be fit to the model to determine its two relevant parameters, the bacterial carrying capacity, *K*, and a factor that characterizes the collapses, *z*. In itself, this is trivial. However, we can also determine *z* from independent, and quite different, data, namely image-based time-series of individual fish. The two measures agree, which provides strong support for the proposed stochastic-collapse-driven model of inter-species competition. Furthermore, the fit of the model to the data reveals that the impact of *Vibrio* on *Aeromonas* populations is twofold: reducing the overall carrying capacity and increasing sensitivity to physical perturbations—the combined effect of the two being much greater than either alone. More generally, our analysis provides evidence that quantitative, data-based models of interactions among species within the gut are possible, and that stochastic, rather than purely deterministic, dynamics can play a major role in shaping the composition of and competition within intestinal bacterial communities. It is interesting to note that recent metagenomic analyses of human intestinal microbiota have uncovered signatures of sudden shifts in species composition, the origins of which remain unknown [15, 42], perhaps indicating stochastic dynamics are widespread in natural intestinal systems.

From an ecological perspective, it is unsurprising that the physical environment and stochastic perturbations influence species abundance; these concepts are mainstays of our understanding of macroscopic multi-species communities [43]. A rich literature describes various stochastic population models and the characteristics, such as extinction probabilities, that emerge from them [44-47]. As shown here, it is likely that such models will in general be useful for providing a conceptual and predictive framework for understanding inter-species bacterial competition. Again mirroring well-established ecological concepts, we can frame our understanding of *Vibrio* and *Aeromonas* dynamics in the intestine as a study of these species’ differential resistance and resilience to environmental perturbations. *Aeromonas* during mono-association is not resistant to disturbances related to intestinal motility, but it is resilient, able to grow to high abundances despite sporadic collapses. *Vibrio*, in contrast, is highly resistant to perturbations; it shows smooth growth unfazed by the environmental perturbations that affect *Aeromonas* (Fig 3). In the presence of *Vibrio*, both the resistance and resilience of *Aeromonas* are compromised, as the magnitude of collapses is greater and the carrying capacity to which to recover is diminished.

While ecological concepts can help us characterize microbial dynamics, data on microbial systems can, conversely, enhance our understanding of ecological theory. The fast generation time and high degree of reproducibility of microbial systems have allowed a variety of tests of ecological models in recent years, illuminating issues such as game-theoretic aspects of cheating [48], early warning indicators of population collapses [49], and the statistical structure of number fluctuations [50]. Although theoretical treatments of population collapses and extinction events are abundant in the ecological literature, real data with which to test them remain sparse [51], in part due to the challenges of performing high-precision field studies. We expect, therefore, that data of the sort presented here, which yield collapse statistics as well as fits to stochastic models, will have utility in contexts far removed from microbiota research.

*Aeromonas* population collapses are well described by stochastic dynamics, but the underlying mechanism by which *Vibrio* compromises resistance and resilience of *Aeromonas* remains to be elucidated. Several possibilities exist, and are the focus of ongoing investigation. *Vibrio* may disrupt the adhesive properties of sessile bacterial communities by secreting mucinases [52], or alter the rheological properties of the intestinal environment [53]. More directly, *Vibrio* may kill *Aeromonas* via secreted factors acting as bacteriocins or contact-mediated killing through the Type VI secretion system [54-56]. Intriguingly, it is unclear whether, in the context of a larger metacommunity composed of many fish in a shared aqueous environment, *Aeromonas* is actually at a competitive disadvantage compared to *Vibrio*. Expulsions of *Aeromonas* could benefit this species by aiding dispersal and subsequent colonization of other hosts. This may, in fact, explain the observation that species of *Vibrio* and *Aeromonas* are both highly represented among conventionally raised zebrafish [31].

The combination of gnotobiotic manipulation and imaging-based analyses can be further elaborated in larval zebrafish, both by increasing the diversity of monitored microbial species and by examining interactions with particular aspects of the host such as its immune system [25]. As illustrated here, we expect that such studies will yield additional insights into the factors that drive the dynamics of complex, natural host-associated microbiota.

## METHODS

### Ethics statement

All experiments with zebrafish were done in accordance with protocols approved by the University of Oregon Institutional Animal Care and Use Committee and following standard protocols [57].

### Gnotobiotic techniques

Wild-type AB or *ret* mutant (*ret1^hu2846^*, ZFIN ID: ZDB-ALT-070315-12) zebrafish were derived GF and colonized with bacterial strains as previously described [19]. Briefly, fertilized eggs from adult mating pairs were harvested and incubated in sterile embryo media (EM) containing 100 µg/ml ampicillin, 5 µg/ml kanamycin, and 250 µg/ml amphotericin B for ~6 hour. Embryos were then washed in EM containing 0.003% sodium hypochlorite followed by EM containing 0.1% polyvinylpyrrolidone–iodine. Sterilized embryos were distributed into T25 tissue culture flasks containing 15 ml sterile EM at a density of one embryo per ml and incubated at 28-30°C prior to bacterial colonization. Embryos were sustained on yolk-derived nutrients and not fed during experiments.

### Bacterial strains

*Aeromonas* (ZOR0001, PRJNA205571) and *Vibrio* (ZWU0020, PRJNA205585) were isolated from the zebrafish intestinal tract and described previously [24]. Fluorescently marked derivatives used in imaging experiments were engineered with an established Tn7 transposon-based approach [58]. Briefly, a cassette containing the constitutively active synthetic promoter Ptac cloned upstream of genes encoding dTomato or superfolder GFP was chromosomally inserted at the *attTn7* locus to generate *Aeromonas attTn7*::*Ptac-dTomato* and *Vibrio attTn7*::*Ptac-sfGFP*. Strains expressing fluorescent proteins did not exhibit overt fitness defects *in vitro* or *in vivo*. Prior to colonization at designated time points, bacterial strains were grown overnight in Luria Broth (LB) shaking at 30°C. Bacterial cultures were prepared for inoculation by pelleting for two minutes at 7,000 x g and washing once in sterile embryo medium (EM). An inoculum of 10^6^ CFU/ml was used across experiments for each bacterial strain and added directly to the water column.

### Culture-based quantification of bacterial populations

Dissection of larval guts was done as described previously [19]. Dissected guts were harvested and placed in a 1.6 ml tube containing 500 µl sterile 0.7% saline and ~100 µl 0.5 mm zirconium oxide beads (Next Advance, Averill Park, NY). Guts were then homogenized using a bullet blender tissue homogenizer (Next Advance, Averill Park, NY) for ~25 seconds on power 4. Lysates were serially plated on tryptic soy agar (TSA) and incubated overnight at 30°C prior to enumeration of CFU and determination of bacterial load. Plots depicting culture-based quantification of bacterial populations show the estimated limit of detection (5 bacteria/gut) as well as limit of quantification (100 bacteria/gut) and represent pooled data from a minimum of two independent experiments.

### Light sheet microscopy

Imaging was performed using a home-built light sheet fluorescence microscope, based on the design of Keller et al. [18] and described in detail elsewhere [21, 22]. Briefly, a laser beam is rapidly scanned with a galvanometer mirror and demagnified to provide a thin sheet of excitation light. An objective lens mounted perpendicular to the sheet captures fluorescence emission from the optical section, and the sample is scanned along the detection coordinate to yield a three-dimensional image. To image the entire extent of the intestine (approximately 1200x300x150 microns) we sequentially image four sub-regions and computationally register the images after acquisition. The entire volume of the intestine is imaged in less than 2 minutes in two colors, with a 1-micron spacing between planes. Unless otherwise indicated in the text, all exposure times are 30 ms with an excitation laser power 5 mW, as measured between the theta-lens and the excitation objective.

### Imaging-based quantification of bacterial populations

The analysis pipeline used to estimate bacterial abundances from light sheet imaging is described in [21]. In brief, we computationally identify both individual bacteria and clusters of bacteria, and estimate the population of each cluster by dividing the total fluorescence intensity by the average intensity of individual bacteria. As necessary, objects that are falsely identified as bacterial clusters are manually removed. For example, in Fig 3A an autofluorescent signal in the intestinal midgut in the *Vibrio* channel was excluded from subsequent quantitative analysis. Additionally, individual time points during time-series are removed if, determined by manual inspection, sample drift or motion of bacterial clusters driven by intestinal motility makes it infeasible to robustly estimate bacterial abundance.

### Identification of population collapse events

Collapses in bacteria populations are objectively identified from time-series of total bacterial abundance, such as those in Fig 3, by defining a collapse as a decrease in population by at least a factor of 10 within one hour. Collapse events with pre-collapse populations of less than 100 bacteria are discarded. These criteria were manually validated by associating each identified collapse with a corresponding ejection of bacteria from the gut observed in series of images.

### Imaging experiments

Sample mounting is done as previously described [21]. Larval zebrafish were removed from culture flasks and anaesthetized using 120 µg/ml tricaine methanesulfonate (Western Chemical, Ferndale, WA). Individual specimens were then briefly immersed in 0.5% agar (maximum temperature: 42° C) and drawn into a glass capillary, which was then mounted onto a sample holder. The agar-embedded specimens were partially extruded from the capillary so that the excitation and emission optical paths did not pass through glass interfaces. The specimen holder can hold up to six samples, all of which are immersed EM maintained at 28°C, with tricaine present as an anaesthetic. All long-term imaging experiments were done overnight, beginning in the late afternoon.

### Measuring bacterial distance to epithelial wall

Individual bacteria were identified using the same algorithms used for quantification of bacterial abundance in the intestine. As we do not have a fluorescent marker for the epithelial wall of the intestine, we use the extent of the autofluorescent mucus in images as an estimate of the location of the epithelial wall. This extent is determined by active contour segmentation using the Chan-Vese algorithm [59], using the implementation provided in MATLAB. A user-defined region is used as the seed for the segmentation of the first frame in the time-series, after which the segmentation of the previous frame is used as the seed for the segmentation of the subsequent frame. We then define the distance of each identified bacterium to the epithelial wall as the minimum distance between the location of the bacterium and the segmented extent of the intestine. Distributions of distances to the epithelial wall are constructed from all video frames and confidence intervals are obtained using bootstrap resampling. A null model of a uniform prediction is obtained by randomly distributing 1000 points for each time point in the region defined by our intestinal segmentation. Confidence intervals are again obtained through bootstrap resampling.

### Measuring intestinal motility

Larval intestinal motility was assessed from images captured using differential interference contrast (DIC) microscopy, performed as previously described [27]. The displacement field from frame to frame in time-series was determined using particle image velocimetry (PIV) algorithms [60], which calculate the motions necessary for regions in one frame to be mapped onto regions in another. We focused our analysis on the frequency and amplitude of these motions, restricting our analysis to components of displacement along the intestinal axis. Fourier spectra of the displacements, averaged over location in the intestine, yielded in all cases a clear peak whose frequency and magnitude are indicative of the characteristic frequency and amplitude of intestinal motility, respectively. This method is described in greater detail in a forthcoming paper.

## ACKNOWLEDGEMENTS

We thank Michael Taormina (University of Oregon) for experimental advice, and Brendan Bohannan (University of Oregon) and Pankaj Mehta (Boston University) for critical readings of the manuscript. For providing the zebrafish knockout allele *ret1-hu2846* we thank the Hubrecht laboratory and the Sanger Institute Zebrafish Mutation Project. This material is based upon work supported by the National Science Foundation under grant no. 1427957. Research reported in this publication was also supported by the National Institutes of Health, including the National Institute of General Medical Sciences (P50GM09891), the National Institute of Child Health and Human Development (P01HD22486), and the National Institute of Allergy and Infectious Diseases (F32AI112094). The content is solely the responsibility of the authors and does not necessarily represent the official views of the National Institutes of Health. We also acknowledge support from Research Corporation and the Gordon and Betty Moore Foundation.

## SUPPORTING INFORMATION

### S1 Text. Stochastic Collapse Model

We describe a simple model of growth and collapse behavior and examine its predictions for population sizes. We also fit the model to experimental data on bacterial abundance.

#### 1 The model

Consider a species with population *N* at time *t* that exhibits logistic growth, with growth rate *r* and carrying capacity *K*:

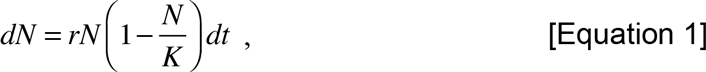

We superimpose on these dynamics events in which the population collapses to a value *f* times its pre-collapse value, where *f* is between 0 and 1, after which it resumes logistic growth. We model the timing of the collapses as a Poisson process: collapses are uncorrelated and stochastic, occurring with a probability per unit time *p*_c_. Formally, one can write this as a stochastic differential equation:

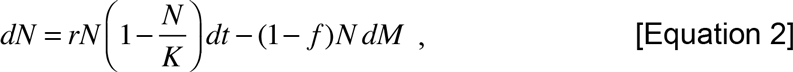

where *dM* is a Poisson process of unit step. (In other words, *dM* =1 with probability *p*_c_*dt*, and *dM =* 0 with probability 1 - *p*_c_*dt*.) *N dM* refers to *N* immediately before the collapse. An illustration of the roles of the parameters *r, K, f*, and *p*_c_ is provided in Figure 5. As noted in the main text, this model is not new; it has been invoked and studied in many ecological contexts [S1]. However, the particular treatment presented here is, to the best of our knowledge, novel, especially with respect to determining relevant parameters for fits to experimental data. We determine statistical properties of the model using numerical simulations. For infinite carrying capacity, these properties can be calculated analytically, but for the biologically relevant case of finite carrying capacity, exact solutions do not at present exist.

#### 2 Simulations

The model described above is simple to simulate by numerical integration, which yields the population *x*_t_ at time *t*. Two typical *x*_t_ are shown in Figure ST1, with parameters as noted in the caption.

**Figure ST1.**
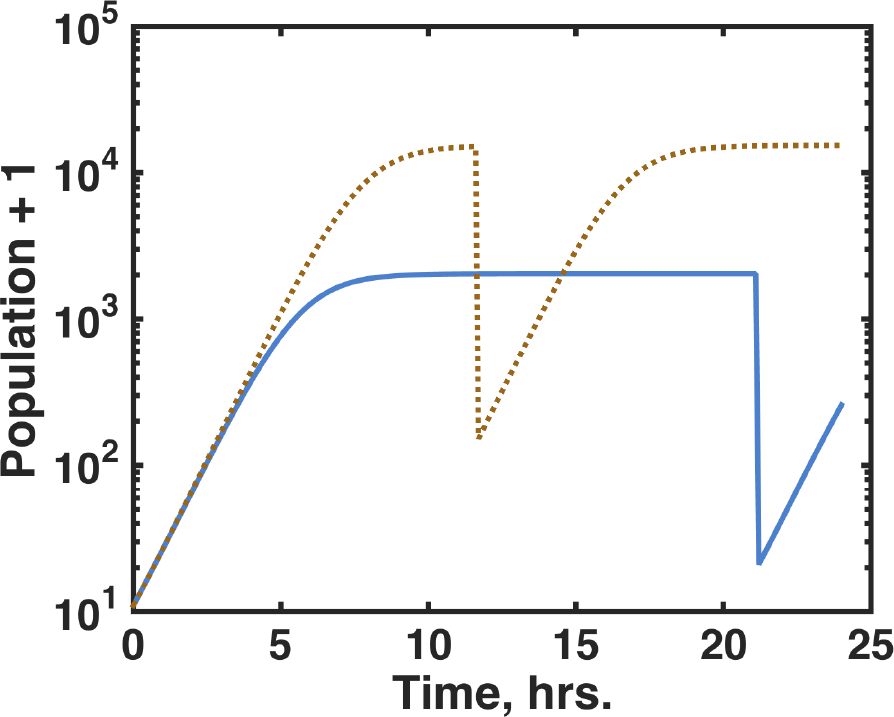
Two simulated populations exhibiting stochastic collapses, with *f =* 10^-2^, *p*_c_ = 0.05 hr.^-1^, *r =* 1 hr.^-1^, and *K* drawn from a log-normal distribution with mean 10^4^ and a standard deviation of half a decade. (We plot the population plus one so that zero values are evident on the logarithmic scale.)

The model has four parameters, *r, K, f*, and *p*_c_, and a boundary condition set by *x*_0_ (the initial population). The value of *x*_0_ is irrelevant for the experimental conditions considered: the populations start from a small value and grow rapidly. In our simulations *x*_0_ is taken to be 10.

The growth rate, *r*, is known from measurements. Moreover, the model dynamics are fairly insensitive to *r*, since the experimental timescales of ~10 hours are considerably larger than the timescale set by the growth rate (1/*r* ~ 1 hour).

The key determinants of the population statistics, therefore, are the collapse properties (*p*_c_ and *f*) and the carrying capacity, *K*. The carrying capacity may exhibit considerable variation between fish. Typically, the final populations of *Aeromonas* in mono-associations are found to be approximately log-normally distributed (Figure ST2), as is commonly the case for species abundances, and so in simulations we draw *K* from log-normal distributions. In other words, log_10_(*K*) for a given simulation is drawn from a Gaussian distribution with some mean value and standard deviation σ_K_, where σ _K_ is typically 0.5, discussed further below. We note that in the absence of collapse (e.g. *p*_c_ = 0 or *f*=1) this model is completely deterministic, and the variance in final bacterial populations between fish is solely due to the variance in *K*.

For particular parameter values, we simulate many instances of the above dynamics (typically 1,000 to 10,000) and examine the statistical properties of the final population, *x*_t_, assessed at *t =* 24 hours. For the values used in Figure ST1 above, for example, the mean and standard deviation of the final *x*_t_ are (6.4 ± 11.5) × 10^3^. The distributions span orders of magnitude, including zero, so it is useful to consider the mean and standard deviation of log_10_(*x*_t_+1), similar to a geometric mean. For these parameters, this gives a mean and standard deviation of log_10_(*x*_t_+1) of 2.8 ± 1.5. We will define *y* as

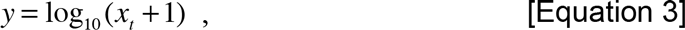

for notational simplicity.

**Figure ST2.**
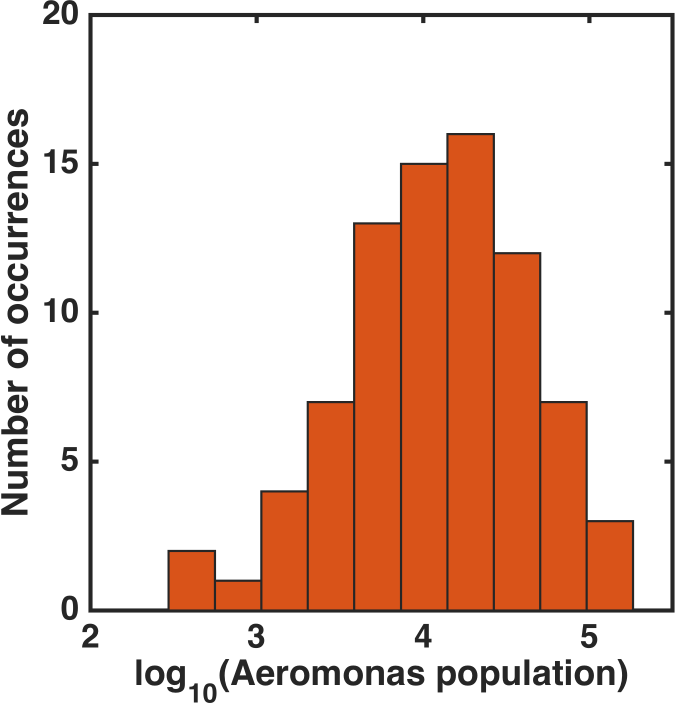
Histogram of the final population of *Aeromonas* mono-associated with larval zebrafish at 4 dpf and assessed at days 5, 6, or 7 dpf by plating of dissected gut contents and counting of colony forming units.

#### 3 Parameters and Fits

##### 3.1 Dependence on *p*_c_ and *f*

We can vary the model parameters to determine the relationship between the mean and the variance of the final population, which will allow direct comparison between our model and measurements of bacterial abundance (e.g. Figure 1). The dependence of the mean and standard deviation (std.) of *y* on *p*_c_ and *f* is plotted in Figure ST3. We can intuitively understand its behavior: for small *p*_c_ or *f* near 1, the properties of *x*_t_ are largely set by the mean and variance of the carrying capacity. However, as *p*_c_ increases (or *f* decreases), the mean of *x*_t_ decreases, because larger collapses are more likely to occur, and the standard deviation of *x*_t_ increases, because the stochastic collapses play a more significant role in the dynamics. For still larger *p*_c_ (or smaller *f*), the final population becomes more uniformly small, because the population is dominated by very frequent collapses and cannot grow appreciably.

Treating *f* as a random, rather than a fixed, parameter has little effect on the behavior of the model. Drawing *f* from a beta distribution, chosen because it is continuous, spans [0, 1], and has two parameters that can be mapped onto a mean and variance, gives the curve shown in Figure ST4. The mean collapse magnitude is chosen over the same range as *f* in Figure ST3, and for each mean *f*, several *f* values are drawn from a beta distribution with standard deviations relative to the mean spanning [0, 0.8]. All the resulting population characteristics are plotted in Figure ST4; the resulting curve is nearly identical to that of Figure ST3.

**Figure ST3.**
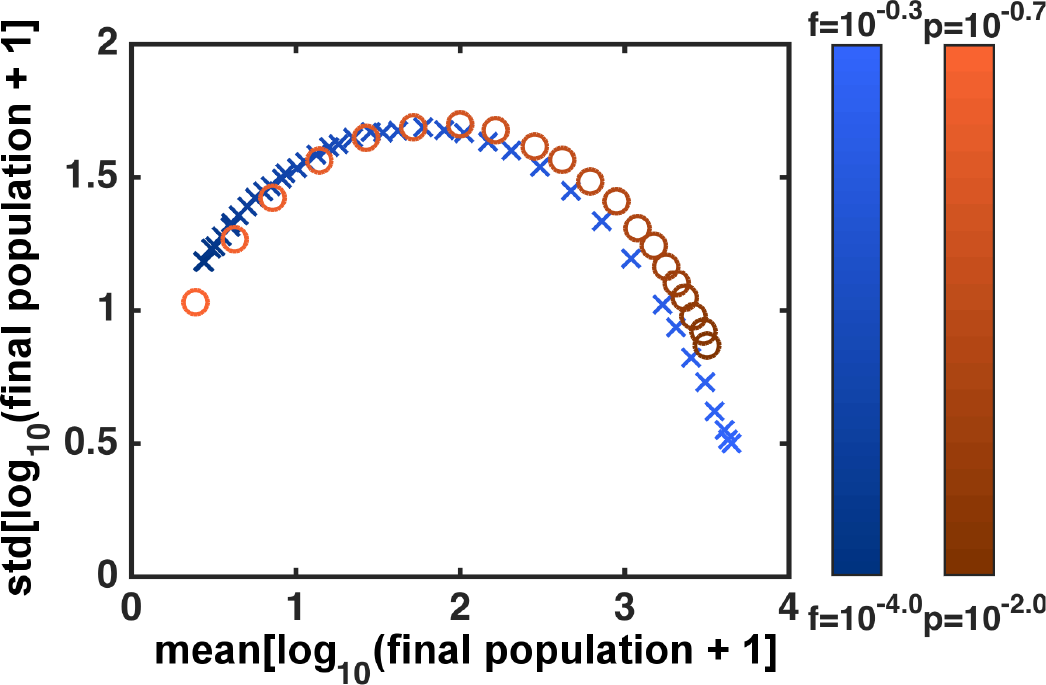
The mean and standard deviation of simulated populations at *t =* 24 hrs., with *r =* 0.8 hr.^-1^ and *K* drawn from a log-normal distribution with mean 10^4^ and a standard deviation of half a decade. Blue crosses: *p*_c_ is fixed at = 0.1 hr.^-1^, and *f* varies between 10^-4^ and 10^-0.3^. Red circles: *f* is fixed at 10^-2^ and *p*_c_ varies between 10^-2^ and 10^-0.7^ hr.^-1^. Each point is calculated from 10,000 simulated runs.

**Figure ST4.**
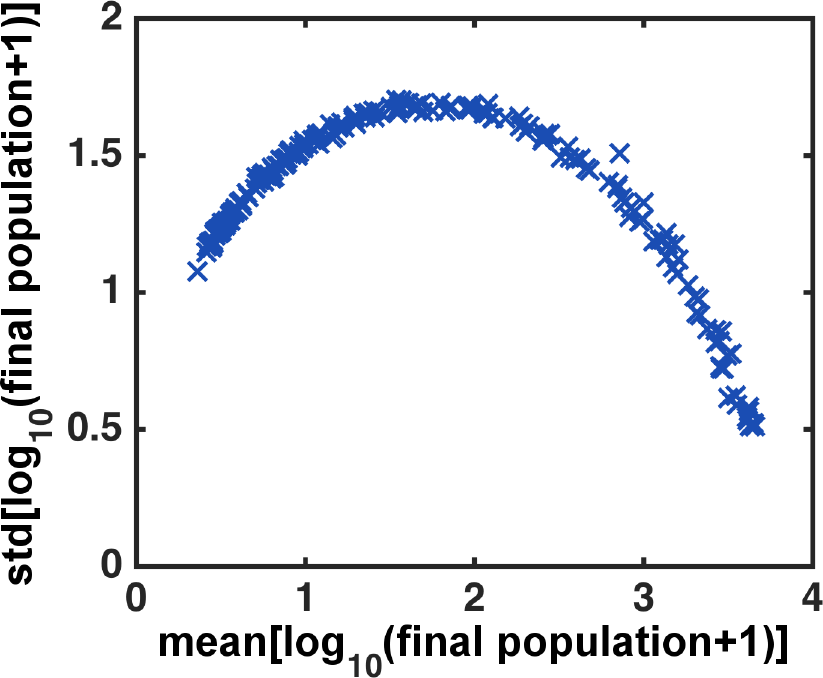
The mean and standard deviation of simulated populations at *t =* 24 hrs., with *r =* 1 hr.^-1^ and *K* drawn from a log-normal distribution with mean 10^4^ and a standard deviation of half a decade. The collapse probability *p*_c_ is fixed at = 0.1 hr.^-1^, and *f* is drawn from a beta distribution with mean between 10^-4^ and 10^-0.3^, and relative standard deviation between 0 and 80%. Each point is calculated from 1,000 simulated runs.

Remarkably, at fixed *K*, nearly identical curves result from varying either *p*_c_ or *f* (Figure ST3), suggesting that at least over the parameter ranges and timescales relevant to our experiments, these two parameters can be subsumed into one effective variable. Considering particular values of mean(*y*) and std(*y*), where *y* is the logarithm of the population as defined above, we can search for the best-fit values of (*p*_c_, *f*), i.e. the parameters that minimize the squared Euclidean distance, χ^2^, between the measured and simulated (mean(*y*), std(*y*)). Using, for concreteness, the values determined from gut dissection and plating experiments of *Aeromonas* abundance 24 hours after challenge by *Vibrio*, namely (mean(*y*), std(*y*)) = (1.68 ± 0.34, 1.50 ± 0.24), we find, as expected, the best-fit contours describe a curve in the (*p*_c_, *f*) space (Figure ST5a). Empirically, we find that this curve is represented by –*p*_c_ log_10_(*f*) ≈ constant (Figure ST5b).

Fitting experimental data to this model of logistic growth with stochastic collapses reduces, therefore, to a two parameter fit to the carrying capacity, *K*, and a parameter describing the collapse properties, denoted as *z*:

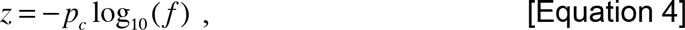

To the best of our knowledge, this effective collapse of the two stochastic parameters into one effective parameter, *z*, has not been previously reported. We do not have a mathematically exact theory for its occurrence, but simply present it as an empirical result from our numerical simulations.

**Figure ST5.**
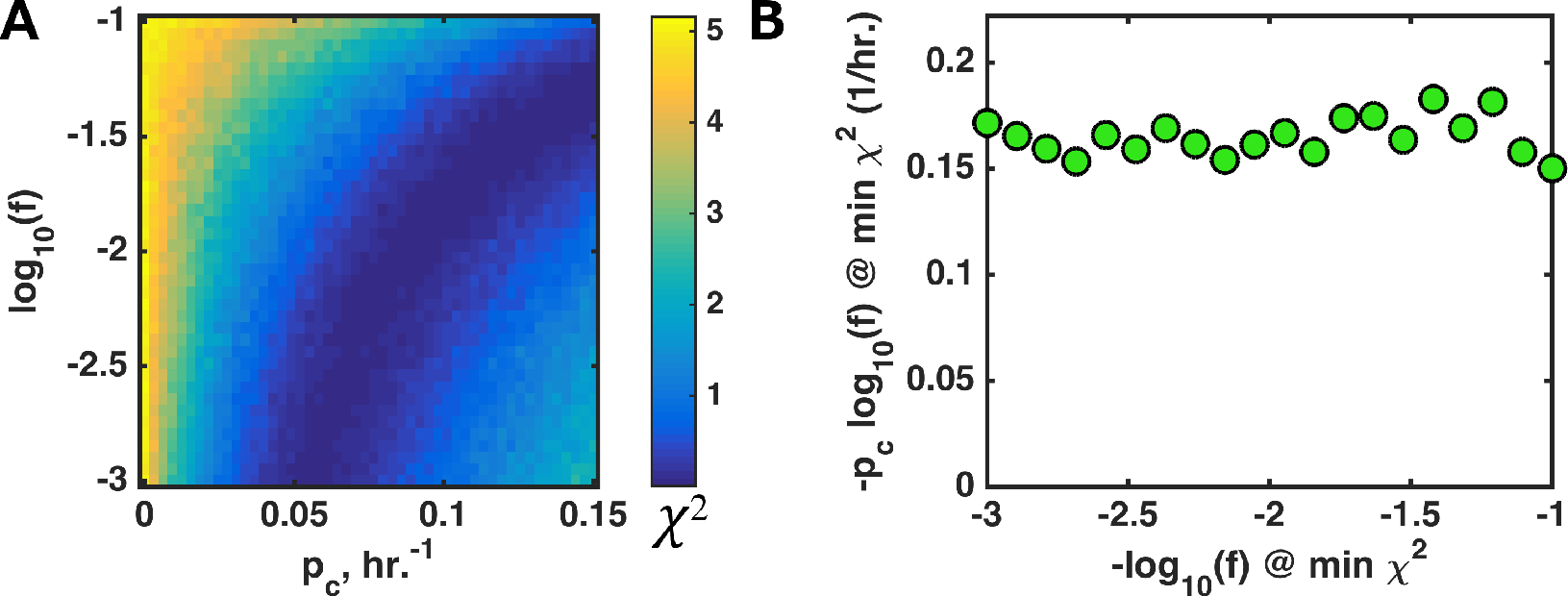
(A) Squared distance, χ^2^, between the measured and simulated (mean(*y*), std(*y*)) for values derived from *Aeromonas* abundance 24 hours after challenge by *Vibrio*, namely (mean(*y*), std(*y*)) = (1.68 ± 0.34, 1.50 ± 0.24), as a function of model parameters *p*_c_ and *f*. The carrying capacity is drawn from a log-normal distribution with mean 10^3.7^ and standard deviation 0.5 decades. At each value of (*p*_c_, *f*), 1000 runs are simulated to determine mean(*y*) and std(*y*). The optimal parameters (darkest blue) sweep out a curve in the parameter space. (B) The optimal *p*_c_ and *f* are related by –*p*_c_ log_10_(*f*) ≈ constant over the range of parameters examined.

##### 3.2 Parameter fits: *Aeromonas* challenged by *Vibrio*

Again using the *Aeromonas* 24-hour post-challenge abundance data (Figure 1), (mean(*y*), std(*y*)) = (1.68 ± 0.34, 1.50 ± 0.24), contours of χ^2^ are shown in Figure ST6. The best-fit parameter values are:

> *z =* -*p*_c_log_10_(*f*) = 0.13 ± 0.05 hr.^-1^,

log_10_(*K*) = 3.2 ± 0.5 In the simulations, *K* is drawn from a log-normal distribution with width 0.5 decades; the fit is insensitive to this width, since the variance in the final population is much greater than 0.5. The uncertainties in *z* and *K* are estimated from simulations spanning the experimental uncertainties in mean(*y*) and std(*y*).

In the main text, we compare these plating-derived measures of the collapse parameters *p*_c_ and *f* to those determined from live imaging.

**Figure ST6.**
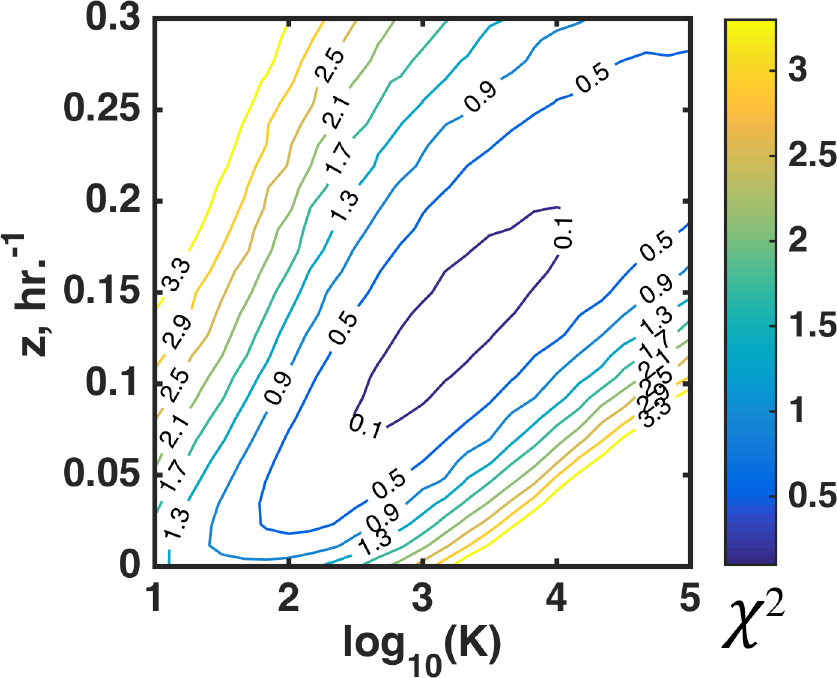
Contours of χ^2^, the distance between simulated (mean(*y*), std(*y*)) and the measured value from di-association experiments (1.68, 1.50), for a range of *z* and *K*. The fit has a clear minimum at *z =* 0.13 hr.^-1^ and log_10_(*K*) = 3.2.

#### 3.3 Parameter fits: *Aeromonas* alone

Similarly, we can determine the parameter values that best match *Aeromonas* mono-association data, (mean(*y*), std(*y*)) = (4.1 ± 0.08, 0.61 ± 0.05), where these values are from plating data at both 5 and 6 days post-fertilization. Because std(*y*) is low, i.e. the data map onto the lower right corner of the curve of Figures ST3-4, it is unclear whether the variance in *y* is due mainly to variance in *K* or to the stochasticity of collapses, and we have no independent measure of the variance in *K*. Considering *K* drawn from log-normal distributions of various widths, we find best-fit values of *z =* -*p*_c_log_10_(*f*) spanning roughly *z =* 0.01 ± 0.01 hr.^-1^, i.e. *z* is poorly constrained. Contours of χ^2^ are shown in Figure ST7. Despite this uncertainty, *K* is well-constrained to be approximately log_10_(*K*) = 4.2 ± 0.1. The significance of this is discussed in the main text.

**Figure ST7.**
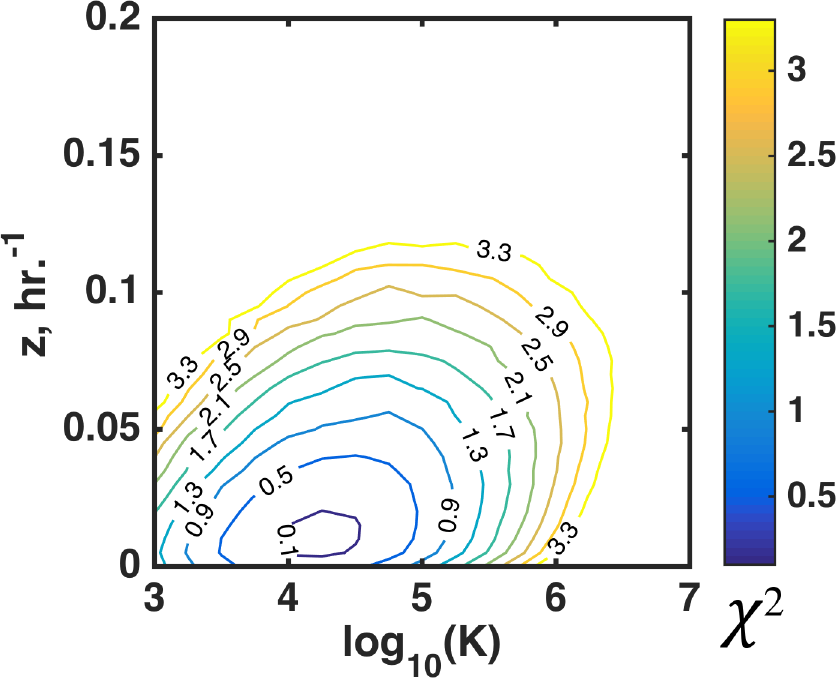
Contours of χ^2^, the squared distance between simulated (mean(*y*), std(*y*)) and the measured value from mono-association experiments (4.1, 0.6), for a range of *z* and *K*, with *K* drawn from log-normal distributions of width 0.1 decades.

## Supporting Figures

**S1 Fig.**
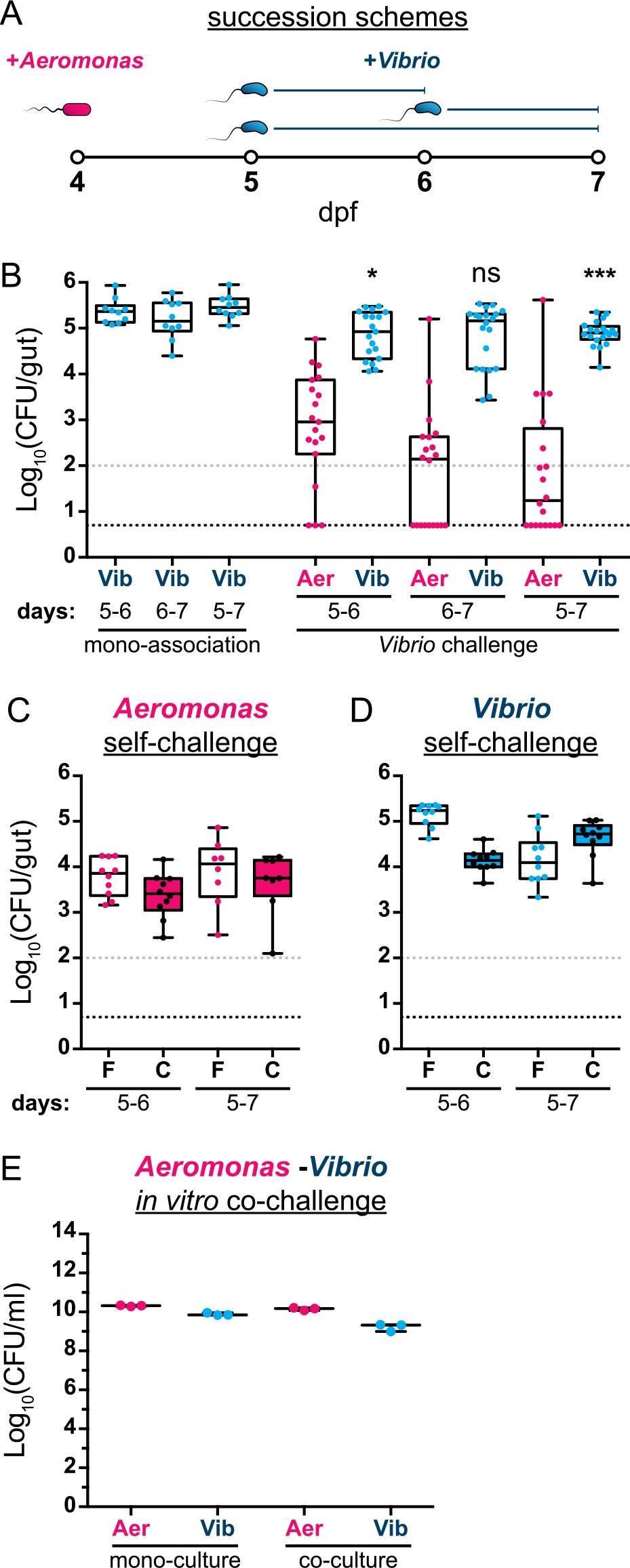
***Aeromonas* and *Vibrio* exhibit an apparent competitive interaction within the larval zebrafish intestine**. (A) Graphical overview of succession schemes used to characterize *Aeromonas*-*Vibrio* interactions. *Aeromonas* is allowed to colonize GF larvae at 4 dpf followed by addition of *Vibrio* to the water column at 5 or 6 dpf for 24 or 48 hours prior to enumeration of abundances by dissection and serial plating. (B, left) *Vibrio* abundances after different mono-association durations and (B, right) *Aeromonas* and *Vibrio* abundances after different *Vibrio* challenge periods. Statistical significance of *Vibrio* abundances after *Vibrio* challenge compared to respective mono-association reference populations (i.e. ‘5-6’ vs. ‘5-6’; ‘6-7’ vs. ‘6-7’; ‘5-7’ vs. ‘5-7’) was determined by an unpaired t-test. *=p<0.05; ***=p<0.0001; ns=not significant; N>10/condition. Founder populations ‘F’ of (C) *Aeromonas* and (D) *Vibrio* were mono-associated with GF larvae on day 4 post-fertilization and challenged by fluorescently marked self populations ‘C’ at 5 dpf for 24hrs (’5-6’) or 48hrs (’5-7’). Dissection and serial plating was done to enumerate founder and challenger populations. Counting of bacterial colonies was done on a fluorescent stereomicroscope. (E) *Aeromonas* and *Vibrio* were inoculated into LB broth either individually or 1:1 and grown overnight with shaking at 30°C prior to enumeration by serial plating. CFU=colony-forming units. Gray and black dashed lines in panels B, C, and D denote limit of quantification and detection, respectively.

**S2 Fig.**
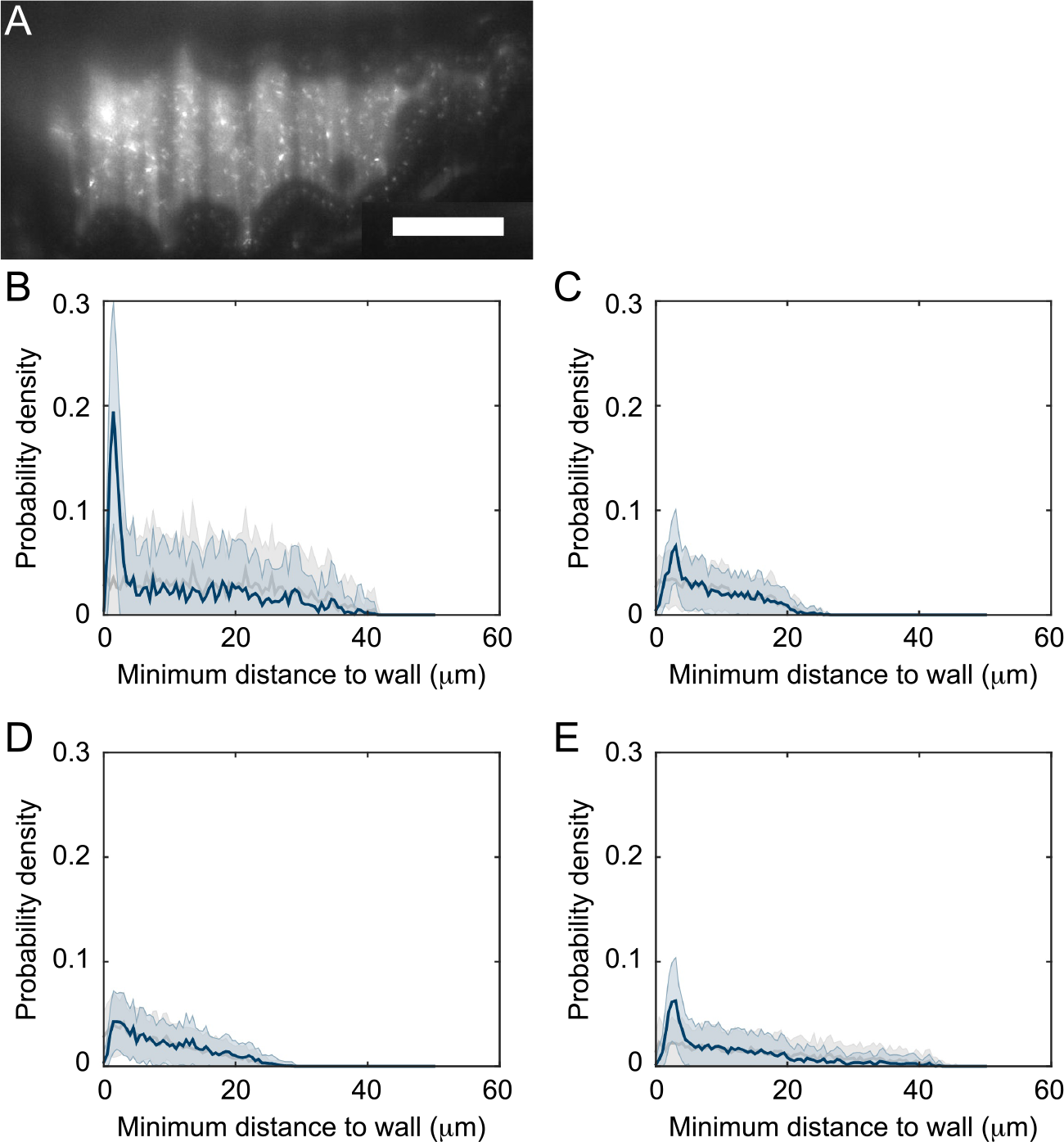
**Space filling properties of *Vibrio* within the zebrafish gut**. (A) Single optical plane of 6 dpf larval zebrafish inoculated at 4 dpf with GFP-labeled *Vibrio*. Scale bar: 50 µm. (B-E) Blue curves: Spatial distribution of bacteria with respect to the approximate extent of the intestinal epithelial wall. Gray curves: prediction from a null model of uniform space filling. Each panel represents an individual fish with panel B being from the same specimen in panel A.

**S3 Fig.**
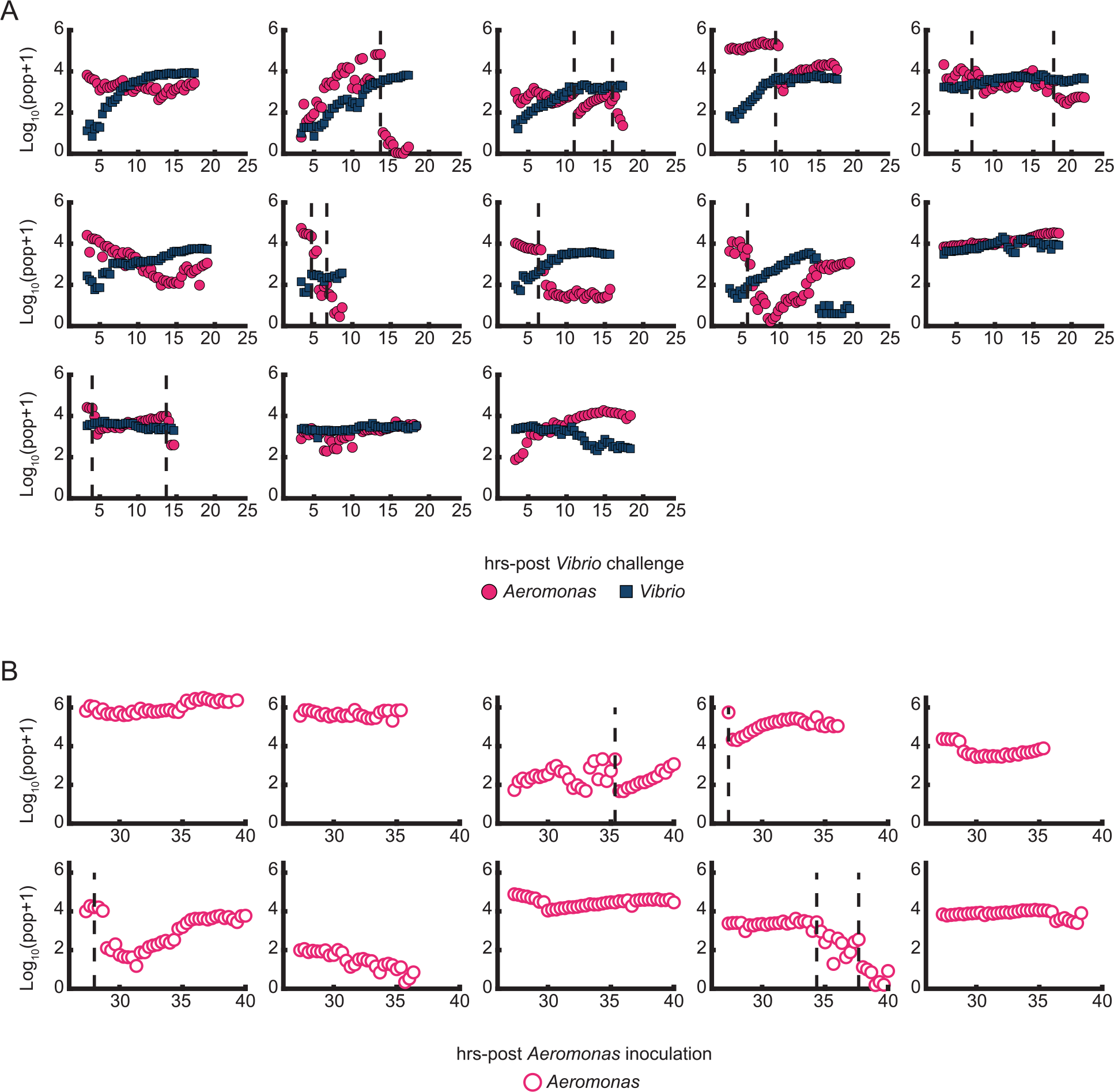
**Collapses of *Aeromonas* populations within the zebrafish gut**. (A) Total bacterial abundance derived from imaging data for *Aeromonas* and *Vibrio* for all imaged fish (*N*=13) initially inoculated for 24 hours with *Aeromonas* and then challenged by *Vibrio*. Plots represent individual larvae and are plotted as a function of time following *Vibrio* inoculation. (B) Total bacterial abundance derived from imaging data for fish inoculated for 24 hours with *Aeromonas* alone (*N*=10). Plots represent individual larvae and are plotted as a function of time following *Aeromonas* inoculation. (A and B) Vertical dashed lines indicate sharp drops of over an order of magnitude within an hour of the *Aeromonas* population.

**S4 Fig.**
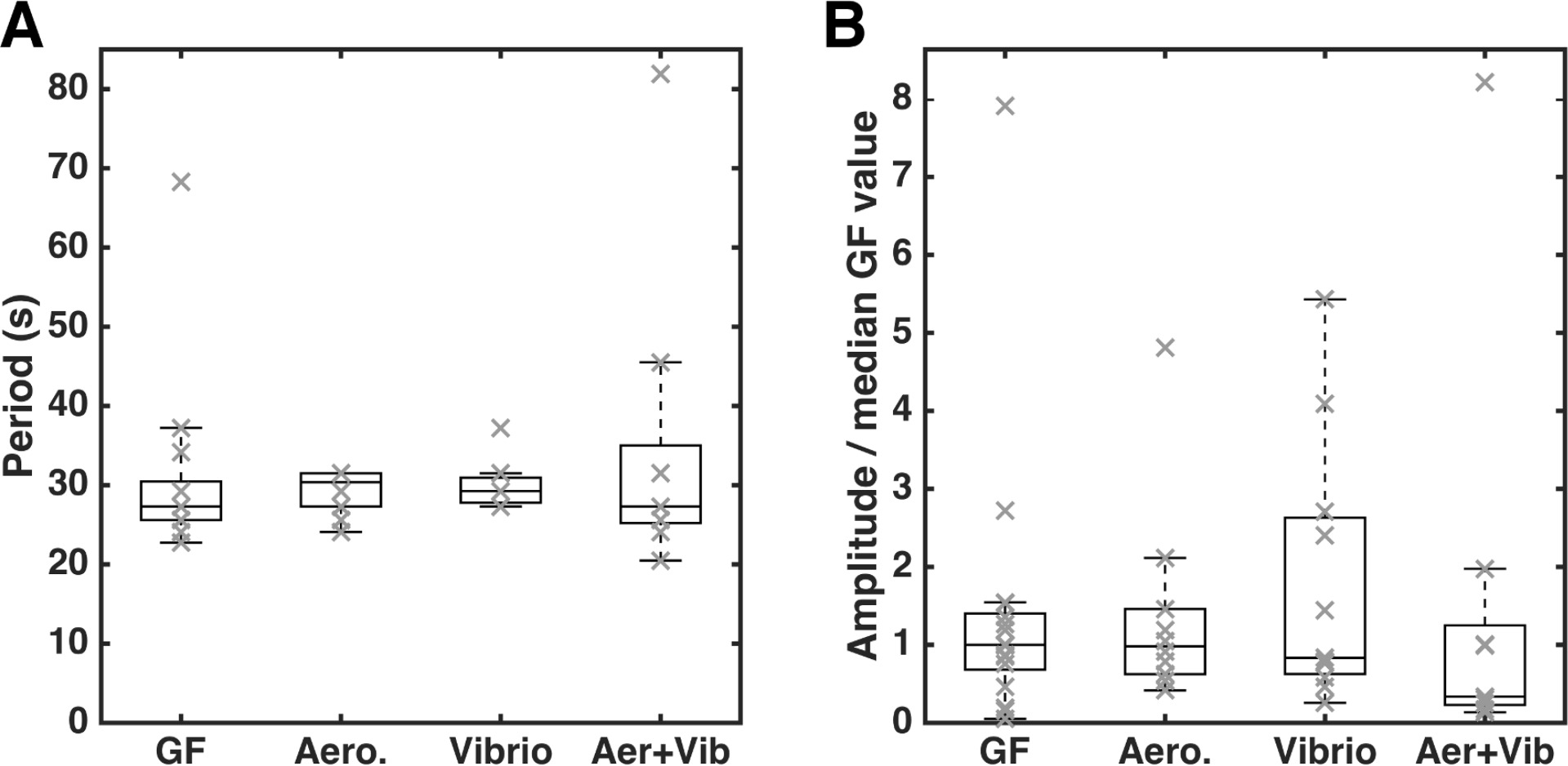
**Characteristics of zebrafish gut motility at 6 dpf for fish with different bacterial colonization histories**. GF = germ-free; Aero = mono-association with *Aeromonas* from 4 dpf; Vibrio = mono-association with *Vibrio* from 4 dpf; Aer+Vib = mono-associated with *Aeromonas* at 4 dpf and challenged with *Vibrio* at 5 dpf. (A) The characteristic period of gut motility, identified as the inverse of the frequency of the peak signal in a Fourier spectrum of gut motion amplitudes, averaged over all positions. All conditions give very similar periodicity of gut motion. (B) The characteristic amplitude of gut motility, identified as magnitude of the peak signal in a Fourier spectrum of gut motion amplitudes. There is considerable variability between fish clutches, and so the amplitudes are normalized by the median of the germ-free fish in each batch. All conditions show large variance, with no significant difference evident between the various conditions. In A and B, gray “X”s are from individual fish; boxes indicate the first to third quartiles, and the horizontal bars in boxes indicates the median value.

**S5 Fig.**
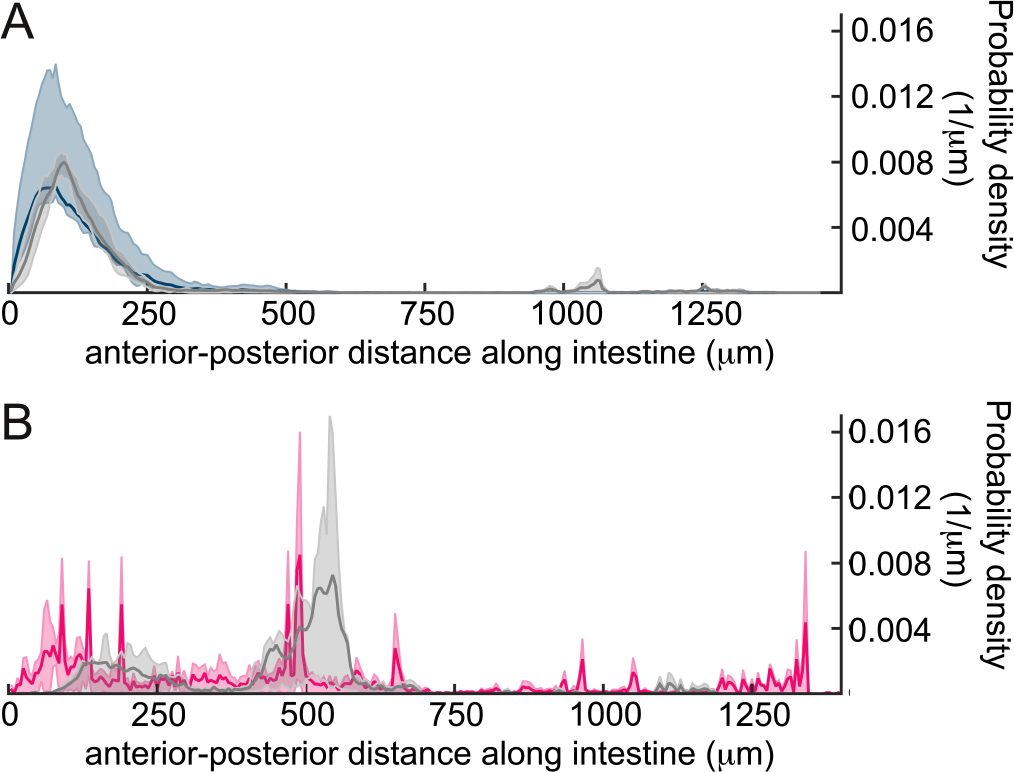
**Spatial distribution of *Aeromonas* and *Vibrio* during mono-association or challenge experiments**. (A) Spatial distribution of *Vibrio*, quantified as the probability density along the gut, for 6 dpf fish mono-associated at 5 dpf with GFP-labeled *Vibrio* (gray) or inoculated at 4 dpf with dTomato-labeled *Aeromonas* and challenged at 5 dpf with GFP-labeled *Vibrio* (blue). (B) Probability density of *Aeromonas* in 6 dpf fish mono-associated at 5 dpf with dTomato-labeled *Aeromonas* (gray) or inoculated at 4 dpf with dTomato-labeled *Aeromonas* and challenged at 5 dpf with GFP-labeled *Vibrio* (magenta). The blue and magenta spatial distributions are drawn from the same fish. *N*=10 for both conditions.

## Supporting Movie Captions

**S1 Movie. Example of the motile and planktonic behavior of *Vibrio* in the zebrafish gut**. Live imaging of a single optical plane in the intestinal midgut of a 6 dpf larval zebrafish inoculated at 4 dpf with GFP-labeled *Vibrio*. Scale bar: 50 µm.

**S2 Movie. Example of *Vibrio* space filling properties**. Three-dimensional scan through the intestinal bulb of a 5 dpf larval zebrafish inoculated at 4 dpf with GFP-labeled *Vibrio*. Scale bar: 50 µm.

**S3 Movie. Example of *Vibrio* resistance to intestinal contractions**. Time-series is of a single optical plane in the intestinal bulb of a 6 dpf larval zebrafish inoculated at 4 dpf with GFP-labeled *Vibrio*. A subpopulation of *Vibrio* can be seen aggregating in the anterior bulb despite repeated intestinal contractions. Scale bar: 50 µm. Movie was recorded at 1 frame per second.

**S4 Movie. Example of the non-motile and clustered behavior of *Aeromonas* in the zebrafish gut**. Live imaging of a single optical plane in the intestinal midgut of a 6 dpf larval zebrafish inoculated at 4 dpf with dTomato-labeled *Aeromonas*. Scale bar: 50 µm.

**S5 Movie. Spatial distribution of *Aeromonas* in the zebrafish gut**. Three-dimensional scan through the intestinal bulb and midgut of a 5 dpf larval zebrafish inoculated at 4 dpf with dTomato-labeled *Aeromonas*. Bacterial clusters, individual bacteria (circled), and autofluorescent signals from intestinal mucus (gray haze) are indicated. Scale bar: 50 µm.

**S6 Movie. Example of an *Aeromonas* collapse event during *Vibrio* challenge**. Time-series is of maximum intensity projections of images taken from the same larval zebrafish shown in Fig 3A. The fish was initially colonized at 4 dpf with *Aeromonas* (magenta), challenged 24 hours later by inoculation with *Vibrio* (cyan), and then imaged every 20 minutes for 14 hours. Times indicate hours post-challenge. The region shown spans about 80% of the intestine, with the anterior on the left. Image contrast in both color channels is enhanced for clarity. Yellow dotted line roughly indicates the lumenal boundary of the intestine; the two bacterial fluorescence channels are overlaid inside this region. Scale bar: 200 µm.

**S7 Movie. Example of *Aeromonas* sensitivity to intestinal contractions**. Time-series is of a single optical plane in the intestinal midgut of a 6 dpf larval zebrafish inoculated at 4 dpf with dTomato-labeled *Aeromonas*. Scale bar: 50 µm. Movie was recorded at 1 frame per second.

**S8 Movie. Example of intestinal motility in a wild-type larval zebrafish**. Differential interference contrast (DIC) microscopy video of intestinal motility in a conventionally raised 6 dpf wild-type larval zebrafish. Scale bar: 50 µm.

**S9 Movie. Example of intestinal motility in a *ret* mutant larval zebrafish**. Differential interference contrast (DIC) microscopy video of intestinal motility in a conventionally raised 6 dpf *ret* mutant larval zebrafish. Scale bar: 50 µm.

